# Ectoparasitism in Polystomatidae (Neodermata, Monogenea): phylogenetic position and mitogenome of *Sphyranura euryceae*, a parasite of the Oklahoma salamander

**DOI:** 10.1101/2022.06.24.497348

**Authors:** Samuel J. Leeming, Christoph Hahn, Stephan Koblmüller, Chris T. McAllister, Maarten P. M. Vanhove, Nikol Kmentová

## Abstract

**Background:** Polystomatidae represents a monogenean group whose representatives infect mainly (semi)-aquatic tetrapods. Sphyranuridae with its single genus (*Sphyranura*) exhibits ectoparasitism on salamander hosts and was traditionally considered a sister-group to Polystomatidae based on the presence of a well-developed opisthaptor yet was distinguished due to the presence of a single pair of haptoral suckers, as opposed to the three pairs present in polystomatids. However, more recent molecular work supported its inclusion within Polystomatidae, at an early diverging, yet unresolved, position in the clade of polystomatids that otherwise exhibit endoparasitism of batrachians. Resolving the position of *Sphyranura* in relation to Polystomatidae is a prerequisite for understanding the factors driving evolution and the shifts between ecto- and endoparasitism in Polystomatidae.

**Methods:** Various staining methods were used to morphologically characterise collected specimens of *Sphyranura*. The mitochondrial genome was assembled from WGS data. Based on a combination of nuclear (*18S, 28S* rRNA) and mitochondrial markers (*cox1, 12S*) we inferred the phylogeny of Polystomatidae using Bayesian inference and maximum likelihood methods.

**Results:** Based on morphological examination and comparison with type material, specimens of *Sphyranura* infecting Oklahoma salamander (*Eurycea tynerensis*) at Greathouse Spring, Arkansas (USA), were identified as *S. euryceae*, a new distributional record for the species. Along with an amended diagnosis of *Sphyranura* we provide the first molecular data for *S. euryceae*. Mitochondrial level comparison reveals instances of tRNA gene rearrangements in polystomatids. Our phylogeny identifies two clades within polystomatids infecting tetrapods, one infecting exclusively batrachians, the other mainly known from chelonians. Although not fully supported, *Sphyranura* appears as the earliest branching lineage within the former.

**Conclusions:** With *Sphyranura* nested within Polystomatidae, we consider Sphyranuridae invalid. *Sphyranura*’s apparent early branching position indicates ectoparasitism is an ancestral trait with endoparasitism having evolved later in the ‘Polbatrach’ clade. However, the reduced number of haptoral suckers in representatives of *Sphyranura* is a derived characteristic potentially resulting from paedomorphic evolution. Whilst there is an indication towards phylogenetic congruence of polystomatids and their batrachian hosts, the same was not true for polystomatid parasites of chelonians with evidence of multiple host switches. Furthermore, geographic distribution of hosts was not found to drive polystomatid phylogeny.

## Introduction

Monogeneans are a globally distributed class of parasitic flatworms of which the vast majority of species are ectoparasites of actinopterygian and chondrichthyan fishes. However, a number of exceptions to this rule are observed where monogeneans of diverse taxa parasitise sarcopterygian hosts. Examples include *Lagarocotyle salamandrae* Kritsky, Hoberg & Aubry, 1993, of the monotypic family Lagarocotylidae, which infects the salamander *Rhyacotriton cascadae* Good & Wake, 1992 [1], *Dactylodiscus latimeris* Kamegai, 1971, a parasite of the coelacanth, representing the monotypic family Neodactylodiscidae Kamegai, 1972 [2], and three members of Lagotrematidae Mañe-Garzon & Gil, 1962 parasitising two species of salamander [3] and a freshwater turtle [4]. The subclass Polystomatoinea Lebedev, 1986 represents a further case of a shift to sarcopterygian hosts in which all but a single species parasitise aquatic and semi-aquatic tetrapod hosts. Furthermore, members of this subclass have also switched from ecto- to endoparasitism in which they typically occupy the urinary bladders of anurans, urodelans and chelonians. Polystomatoinea was long considered to consist of two families, Sphyranuridae Poche, 1925 and Polystomatidae Gamble, 1896 [1,5], traditionally differentiated from one another based on haptoral morphology [6,7]. The former family currently comprises four species in *Sphyranura* Wright, 1879. These are restricted to North America where they infect the gills of salamanders. The latter comprises around 200 species across 26 genera with a global distribution and infecting diverse host taxa [8–10]. Based on structural similarities of the suckers, eggs, and caudal hooks members of *Sphyranura* were originally placed within Polystomatidae [11] but later removed and assigned to Sphyranuridae [12] on the basis that members of *Sphyranura* have a single pair of haptoral suckers in contrast to three pairs found in other polystomatids [13]. *Sphyranura* consists of: *S. osleri* Wright, 1879, *S. oligorchis* Alvey, 1933, *S. polyorchis* Alvey, 1936 and *S. euryceae* Hughes & Moore, 1943. It has been argued, however, that *S. polyorchis* cannot be justified as a separate species from *S. osleri* on the basis of minor morphological differences [14]. While *S. osleri, S. oligorchis* and *S. polyorchis* parasitise the Common mudpuppy (*Necturus maculosus* Rafinesque, 1818) with records of *S. oligorchis* also parasitising the Red River mudpuppy (*Necturus louisianensis* Viosca, 1938) [15], *S. euryceae* is a parasite of the Oklahoma salamander (*Eurycea tynerensis* Moore & Hughes, 1939) [6,16] but has more recently been observed to also parasitise the Cave salamander (*Eurycea lucifuga* Rafinesque, 1822) [6] and Grotto salamander (*Eurycea spelaea* Stejneger, 1892) [17]. In general, there is a scarcity of records of representatives of *Sphyranura* and relatively little knowledge about the genus besides morphology and principal host distribution. However, given the intervening decades since Hughes & Moore’s [18] description of *S. euryceae* advances in staining procedures and microscopy allow for a more detailed morphological examination. Thus, descriptions of representatives of *Sphyranura* often lack some of the morphological information available for more recently studied monogeneans.

Sinnappah et al. [19] inferred a phylogeny of Polystomatoinea based on partial sequences of the *18S* rDNA marker, which placed *Sphyranura* within Polystomatidae. These authors further proposed that the morphological differences between *Sphyranura* and Polystomatidae as described above are the result of an evolutionary retention of juvenile characters in adults within *Sphyranura* [19]. However, this phylogeny only included seven representatives of Polystomatidae and a single representative of *Sphyranura*. Furthermore, the position of *Sphyranura* was not well supported. Subsequent work, also based on partial *18S* rDNA sequences, split Polystomatidae into two lineages: one parasitising exclusively amphibians, the other parasitising mainly chelonians. This phylogeny also supported *Sphyranura* as being nested within the lineage of anuran polystomatids, its exact relationships, however, remained unresolved [20]. More recently, Héritier et al. [21] inferred the phylogeny of Polystomatidae based on the complete *18S* rDNA sequence, a partial *28S* rDNA sequence and two partial mitochondrial genes *cox1* and *12S* rDNA, which supported the division of Polystomatidae into the ‘Polbatrach’ and ‘Polchelon’ lineages with *Concinnocotyla australensis* (Reichenbach-Klinke, 1966), a parasite of the Australian lungfish (*Neoceratodus forsteri* (Krefft, 1870)), branching off prior to this split. The former lineage includes all polystomatids of batrachian hosts (Caudata and Anura) whilst the latter includes all polystomatids of chelonian hosts as well as *Nanopolystoma tinsleyi* du Preez, Badets & Verneau, 2014 of a caecilian host (*Typhlonectes compressicauda* Duméril & Bibron, 1841) and *Oculotrema hippopotami* Stunkard, 1924 of the common hippopotamus (*Hippopotamus amphibius* L., 1758). Furthermore, this phylogeny suggested that *Sphyranura* is an early branching lineage within the ‘Polbatrach’ polystomatids [21]. This phylogeny therefore supported the hypothesis of an origin of Polystomatidae prior to the colonisation of terrestrial environments by tetrapods followed by host-parasite coevolution as different tetrapod lineages diverged [22].

Due to the general paucity of *Sphyranura* records, the genus is notably underrepresented in phylogenetic studies. In this vein all three phylogenetic studies mentioned above were limited, as molecular data representing *Sphyranura* were restricted to *S. oligorchis*. The phylogenetic position of *Sphyranura* in relation to Polystomatidae could therefore be better determined with the inclusion of more representatives of this genus. Specimens of *Sphyranura* sp. were collected from *E. tynerensis* and identified to species level based on morphological examination.

We aim to produce both an amended diagnosis of the genus as well as to provide a clearer picture of the phylogenetic position of *Sphyranura* in relation to Polystomatidae. Further, as the mitogenome of only a single polystomatid is currently available, the level of variation in gene order or presence of non-coding regions, which widely varies between other parasitic flatworm taxa can be assessed. Furthermore, a clarification of the phylogenetic position of *Sphyranura* could shed new insights on the colonisation of salamanders by monogeneans. For instance, a sister group relationship with the other known polystomatid parasitising a salamander (*Pseudopolystoma dendriticum* Osaki, 1948), would indicate the phylogeny of polystomatids may be influenced by that of their tetrapod hosts. An alternative evolutionary hypothesis is that host switches of polystomatids are geographically constrained. In other words, the generally poor dispersal ability of batrachians and chelonians may limit the ability of polystomatids to encounter and colonise new hosts other than those in geographic proximity. Under such a scenario one would expect *Sphyranura* to form a sister-group with other North American polystomatids, regardless of host phylogeny. Unlike the majority of polystomatids, members of *Sphyranura* exhibit ectoparasitism, infecting the skin and gills of their host. The other polystomatid parasitising skin and gills is *C. australensis*, which in the phylogeny of Héritier et al. [21] branches prior to the split between the ‘Polchelon’ and ‘Polbatrach’ clades. The clarification of the phylogenetic position of *Sphyranura* would therefore shed light on the evolutionary transition between ecto- and endoparasitism in polystomatids. Were *Sphyranura* to either branch off prior to the ‘Polbatrach’ - ‘Polchelon’ split, or indeed not be nested within Polystomatidae at all, the transition from ecto- to endoparasitism could be attributed to a single event. On the other hand, were *Sphyranura* to be nested within the ‘Polbatrach’ clade as suggested by Héritier et al. [21] but at a late branching position, this would point to a secondary shift back to ectoparasitism.

## Methods

### Sampling

Over three sampling occasions between November 2019 and November 2020, specimens of *E. tynerensis* were collected with an aquatic dipnet at Greathouse Spring in Tontitown, Benton County, Arkansas, USA (Coordinates 36° 8’ 11.1192’’ N, −94° 12’ 10.0764’’ W). Specimens were placed in habitat water and examined for ectoparasites within 24 hours. Salamanders were killed with an overdose of a concentrated solution of tricaine methanesulfonate and their gills and body examined under a stereomicroscope. When monogeneans were observed on gills, they were removed and relaxed in hot tap water and stored in either 10% neutral-buffered formalin or 98% molecular grade ethanol.

### Staining procedure

Seven adult individuals and two larvae used for morphological analysis were selected from those preserved in 10% neutral-buffered formalin. These were then stained with various media and mounted on standard microscope slides to be morphologically characterised. The staining procedure included the following steps: Individual worms were first placed in a solution of 70% ethanol to be dehydrated before being overstained using a 1:1 mixture of acetocarmine (or Schneider-acetocarmine in the case of specimens 4, 6 and larva 1) and 70% ethanol (>12 hours). The ethanol-acetocarmine mix was then gradually washed out using acid alcohol until internal structures such as testes, ovaries and vesicles were visible under a binocular microscope. At this point the process was halted by washing in distilled water for 5 minutes to remove excess acetocarmine. Specimens 1 and 3 were then stained with Astra blue for 40 minutes before being washed twice in distilled water to wash out residual Astra blue [23]. This step was skipped for specimens 2, 4, 5, 6, 7 and the two larvae. After this, specimens were dehydrated through a series of increasing ethanol concentrations (5 minutes at 70%, 5 minutes at 80%, 15 minutes at 96%, 5 minutes at 100%) and carboxyl was added. Xylene was then added to clear the specimens and they were mounted on a slide using Canada balsam, ensuring that the specimens were lying flat when the cover slip was added. The slides were then weighted to ensure specimens remained flat and given two weeks on a radiator to dry out. The attachment structures of two individuals were placed on a slide in a drop of water that was subsequently replaced by Hoyer’s medium and covered with a cover slip that was sealed with glyceel [24].

### Morphological characterisation

The morphological part of the study was done using Leica DM 2500 LED microscopes (Leica Microsystems, Wetzlar, Germany) and the software LasX v3.6.0 using Differential Interference Contrast (DIC) and Phase Contrast, where necessary, to gain optimal view of individual anatomical features. In total, 35 morphological characters including hard and soft parts were measured following the terminology of [22]. A comparison of the new specimens with existing type material belonging to *Sphyranura* provided by the American Museum of Natural History was undertaken to further support the species identification of these specimens with re-measurements of type material being undertaken where necessary and possible. The material included two specimens of *S. osleri* (Accession numbers AMNH 1427.1 and AMNH 1427.2), one specimen of *S. polyorchis* (Accession number AMNH 1431) and three specimens of *S. oligorchis* (Accession numbers AMNH 1432.1, AMNH 1432.2 and AMNH 1432.3). Pictures of the type material of *S. oligorchis* (AMNH 1432.1) are provided in Supplementary materials S1. Parasite voucher material collected as a part of the present study was deposited in the collection of the American Museum of Natural History under accession numbers xx-xx and Hasselt University under accession numbers xx-xx.

### Molecular Methods

Genomic DNA was extracted using the Quick-DNA™ Miniprep Plus Kit (Zymo Research) following the manufacturer’s instructions with minor modifications, initial incubation overnight, and elution in 2 × 50 µL after 10 min incubation at room temperature each. DNA was then quantified with a Qubit fluorometer (dsDNA HS assay). The DNA concentration of the individual extracts measured between 0.665 and 1.34ng/µl. The complete *12S*, and partial *28S* and *18S* rRNA genes of four specimens were then amplified and sequenced. Primers used for amplification and sequencing of each gene were selected based on previous work [21,25] and were as follows: *18S*: IR5/L7, *12S*: 12SpolF1/12SpolR9, for the *28S* two overlapping fragments of unequal length were sequenced. LSU5/IR14 primers were used for larger of these and IF15/LSU3 for the smaller. The reactions were performed in a total volume of 11µl, including 7.05µl water, 1.0µl buffer (BioTherm 10x PCR Buffer), 0.35µl dNTPs (10mM), 0.25µl forward and reverse primers (1pM), 0.3µl *Taq* polymerase (Supratherm 5 units/µL) and 2.0µl DNA template. The amplification cycle consisted of a step of 3 minutes at 95°C for initial denaturation; 45 cycles of 30 seconds at 95°C for denaturation, 30 seconds at 50°C for annealing and 1 minute at 72°C for elongation; one final step of 7 minutes at 72°C for terminal elongation. The PCR products were visualised on agarose gels in order to verify the success of PCR amplification before sequencing. The PCR products were purified by adding a mixture of 0.5µl ExoSAP (ExoSAP-IT: Amersham Biosciences) and 1.2µl water to each and incubating in a PCR machine for 45 min at 37°C followed by 15 min at 80°C. The sequencing reaction was run using a cycle beginning with a single step of initial denaturation for 3 minutes at 94°C; 35 cycles of 30 seconds at 94°C, 30 seconds at 50°C and 3 minutes at 60°C; one final step of 7 minutes at 60°C. Sequencing products were purified with SephadexTM G-50 (GE Healthcare) and sequenced on an ABI 3130xl capillary sequencer (Applied Biosystems). All newly generated sequences have been deposited on GenBank (see Table 1). DNA extracts of two specimens (SPY1 and SPY2) were sent for whole genome sequencing to commercial sequencing centres. For SPY1 library preparation (Nextera XT, 550 bp insert size) was performed by Macrogen Inc. (Seoul, Korea). For SPY2 library preparation (NEBNext® Ultra IIDNA Library Prep Kit, 550 bp insert size) was done by Novogene (UK). Libraries were sequenced on NovaSeq6000 systems (2×150bp) at the respective centres. Raw read data was first trimmed using Trimmomatic v.0.38 [26] and the following parameters: a minimum length of 40bp, a window size of 5 and required quality per window of 15 and a leading and trailing quality of 3. For both specimens, a subsample of 10,000,000 trimmed reads were randomly selected using seqtk v.1.3 [27] with the seed 553353 and fed into the assembly process. A successful assembly of SPY2 was retrieved using GetOrganelle v. 1.7.1 [28]. A full-length mitochondrial genome of SPY1 could not be recovered using GetOrganelle and so this sample was assembled via MITObim, using the successful SPY2 assembly as a reference. Annotation was then performed via MITOS v.1.0.5 [29] using the genetic code 09 (Echinoderm/Flatworm Mitochondrial). Upon initial visual inspection and comparison of protein-coding genes with those of other monogeneans it became apparent that there were errors in the start and end positions of many protein coding genes given by MITOS v.1.0.5. The assembly was subsequently submitted to MITOS2 via webserver [30]. Start and end positions of protein coding genes as well as start/stop codons were then decided based on visual comparison of the results of MITOS v.1.0.5, MITOS2 and 5 other monogenean species (*D. hangzhouensis*: JQ038227.1, *Neomazocraes dorosomatis*: JQ038229.1, *Microcotyle caudata*: MT180126.1, *Polylabroides guangdongensis*: JQ038230.1, *Neoheterobothrium hirame*: MN984338.1) selected based on the highest percentage identity to the mitogenome of SPY2 when performing a BLAST search. Raw Illumina reads contributing to the mitochondrial genome assemblies were submitted to SRA (accession: xxx) under BioProject accession xxx.

**Table 1.**
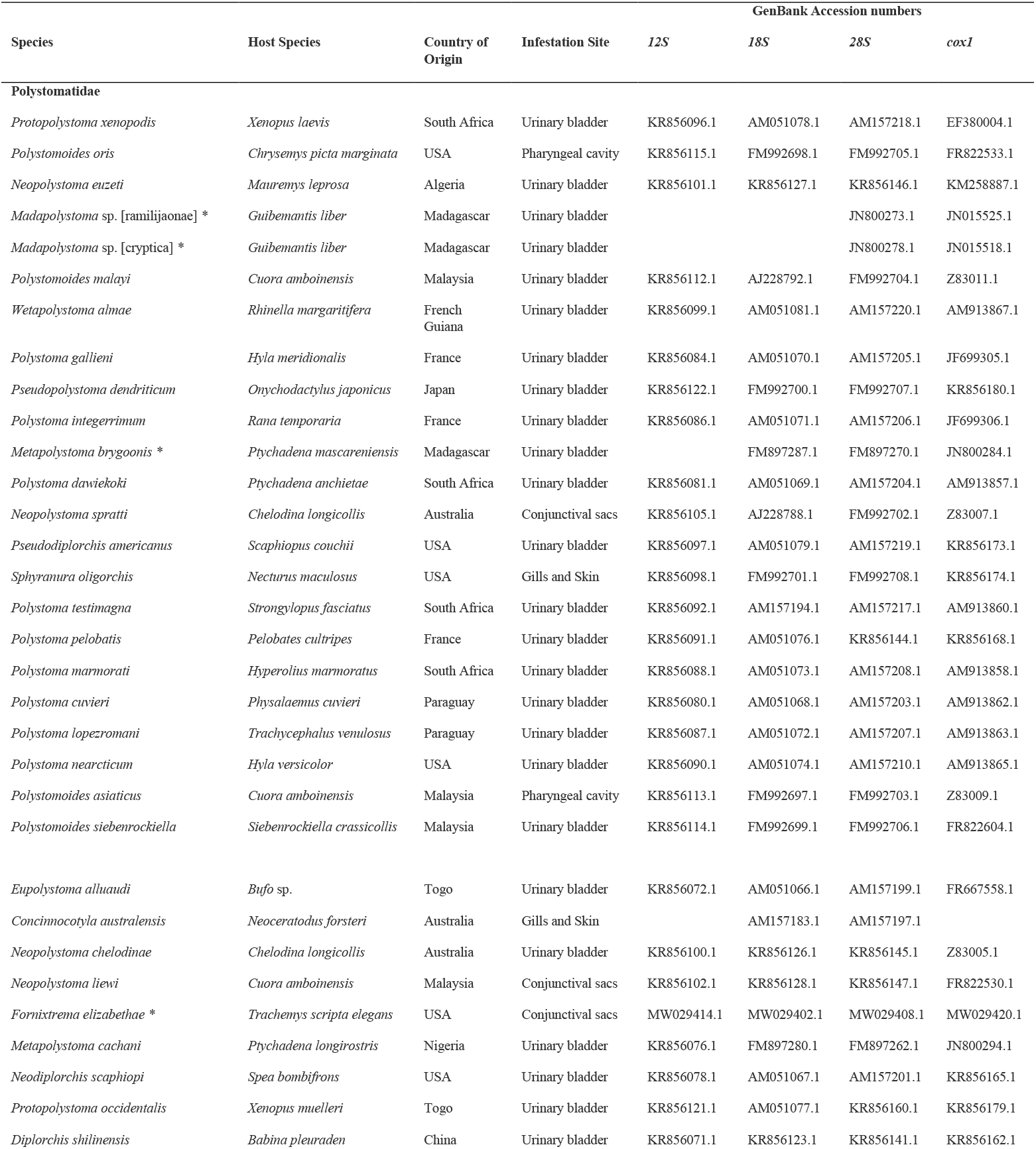
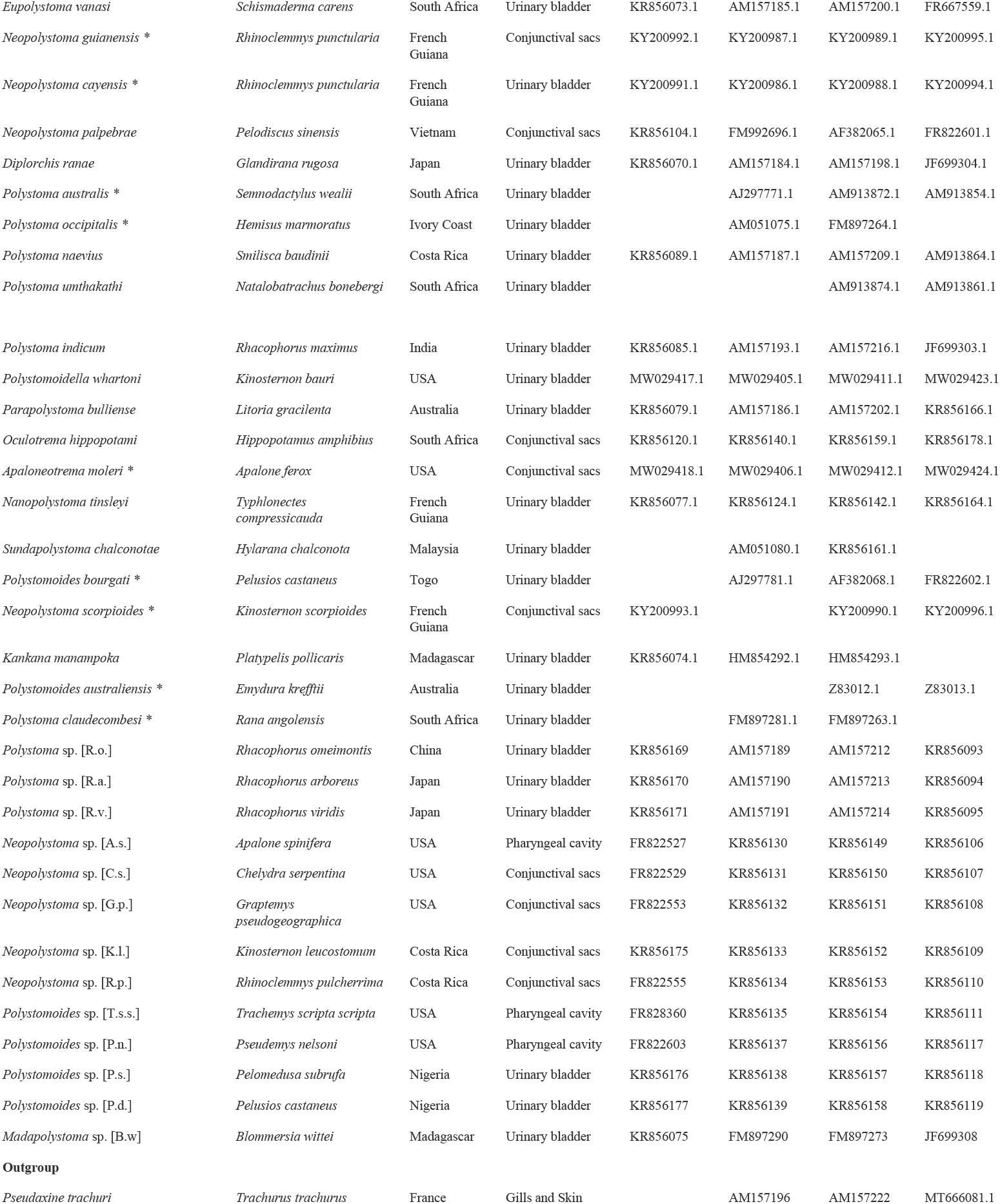
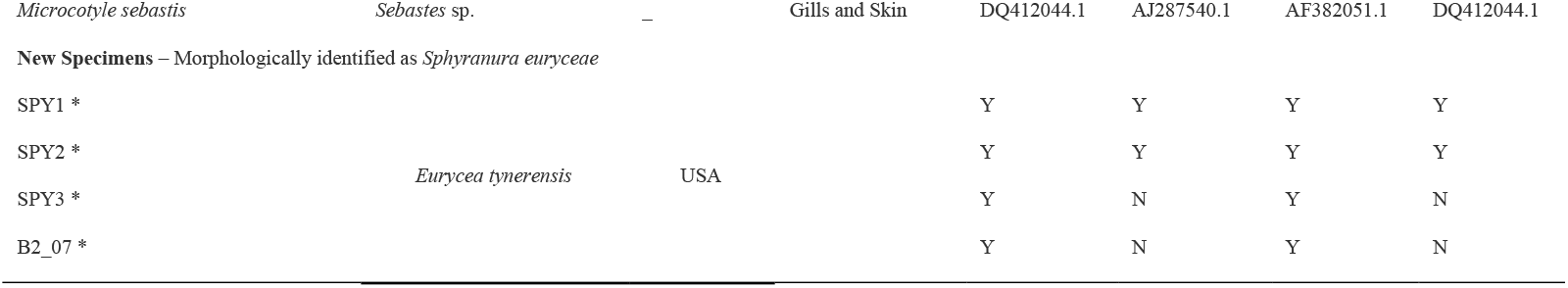
List of parasite taxa and their respective host species, country of origin and GenBank accession numbers of the markers used to infer phylogeny. Taxa marked with * were not included in the phylogeny of Héritier et al. [21].

In addition to MITOS v.1.0.5 the coordinates and secondary structure of mitochondrial tRNA genes was confirmed using ARWEN v.1.2 [31]. In cases where the coordinates given by MITOS v.1.0.5 did not match those of ARWEN v.1.2, those provided by ARWEN v.1.2 were used, provided a 6-7bp acceptor stem was present. The *cox1* and *12S* sequences for the samples SPY1 and SPY2 were retrieved from the mitochondrial genomes based on the annotation results from MITOS2. The mitochondrial genome of SPY1 was compared with that of *Diplorchis hangzhouensis* (Accession: JQ038227.1), the only polystomatid species of which the mitochondrial genome is available, albeit unpublished. Two mitochondrial genomes of *S. euryceae* (SPY1 and SPY2) were deposited on NCBI Genbank under the accession numbers XXX.

Whilst only partial *18S* sequences were retrieved via Sanger sequencing, the complete *18S* sequence could be extracted from WGS data for the samples SPY1 and SPY2. This was done first using Mirabait v.4.9.6 [32] to bait the trimmed reads with the partial *18S* sequences from the respective samples. The baited reads were then interleaved using BBmap v.38.90 [33] and the interleaved reads were assembled via MITObim v.1.9.1 [34] using the partial *18S* sequence as a seed. Barrnap (BAsic Rapid Ribosomal RNA Predictor) v.0.9 [35] was then employed to predict the location of the *18S* sequence within the assembled data. In the case that the newly assembled *18S* sequences were still not equal to the length of the complete *18S* sequences the newly assembled *18S* sequence was used to bait the trimmed WGS reads, and the process was repeated until the complete length of the *18S* sequence was obtained. The same process was employed to obtain the complete length of the *28S* sequences for the samples SPY1 and SPY2.

In addition to sequences obtained from the new specimens, sequences representing a further 65 polystomatid taxa and 2 non-polystomatid monogeneans were accessed via NCBI GenBank. Taxa included in this phylogenetic analysis were selected based on the availability of sequences on NCBI Genbank. A given taxon was included in the analysis on the basis that at least two of the four markers (*12S, 18S, 28S* and *cox1*) were present. Partial sequences were included provided they overlap at least in part with the sequences of all other taxa for which sequence data of a given marker was included. In addition to the 55 polystomatid taxa included in the analysis of Héritier et al. [21], sequences from a further 10 polystomatids were included in addition to the new specimens of *Sphyranura*. Species of Gastrocotylidae (*Pseudaxine trachuri* Parona & Perugia, 1890) and Microcotylidae (*Microcotyle sebastis* Goto, 1894) were selected as an outgroup. Accession numbers of these sequences as well as information on the respective host species, country of origin and site of infection are provided in Table 1.

### Phylogenetic analysis

The four sequence sets were aligned per marker using MAFFT v.7.310 [36] and trimmed using TrimAl v.1.2 “strict mode’’ [37]. Selection of trimming criteria was based upon visual inspection in AliView v.1.28 [38]. The four separate alignments were then concatenated into a single alignment using the script concat.py v.0.21 (https://github.com/reslp/concat). The best fitting partitioning schemes for the three ribosomal sequences as well as the three codon positions of the *cox1* gene were selected by PartitionFinder2 [39] using the “greedy search” algorithm. PartitionFinder2 selected a GTR+I+G model for the three ribosomal subsets as well as the third codon position of *cox1* and a TIM+I+G model for the first two codon positions of *cox1* for IQ-TREE and a GTR+I+G model for all subsets for MrBayes [40]. The Maximum Likelihood tree was run using IQ-TREE v.1.6.12 [41] and a Bayesian tree using MrBayes v.3.2.7 [40] with eight chains running for 70 million generations and sampled every 100 cycles. Phylogenetic trees were visualised using the web-based tool ITOL (Interactive Tree Of Life) [42]. Phylogenetic trees and DNA alignments are openly available in Mendeley Data at https://mendeley.data.com, xxx.

## Results

### Taxonomic account

**Family Polystomatidae Gamble, 1896**

**Genus *Sphyranura* Poche, 1925**

### Amended diagnosis

Body elongated with the greatest body width found approximately half to two-thirds of the distance between the haptor and the oral sucker. Body width (measured at widest point) 17 – 45% of body length with variation between both species and individuals (Table 2). Oral suckers either terminal or subterminal varying in width from 105 – 300μm. Single pair of roughly circular haptoral suckers and of anchors, seven pairs of marginal and one pair of acetabular hooks situated at the basal end of the body. Interior haptoral sucker width accounts for 61 – 68% of haptor width. Haptor length accounts for 14 – 19% of body length and haptor width accounts for 26 – 110% of body width. Vitellaria arranged laterally on both sides of the body extending from the region of the uterus to the peduncle, accounting approximately for two thirds of the body length. Testes intercaecal, arranged either in a single central row or bunched together along the central line of the body. Two excretory vesicles at the level of genital bulb with dorsal openings. Intestinal bifurcation just posterior to pharynx, fused at the level of peduncle. Genital bulb glandular, armed with distally pointed spines. Exhibit ectoparasitism, occupying the skin and gills of caudate hosts (*Eurycea tynerensis, E. lucifuga, E. spelaea, Necturus maculosus & N. louisianensis*).

**Table 2.**
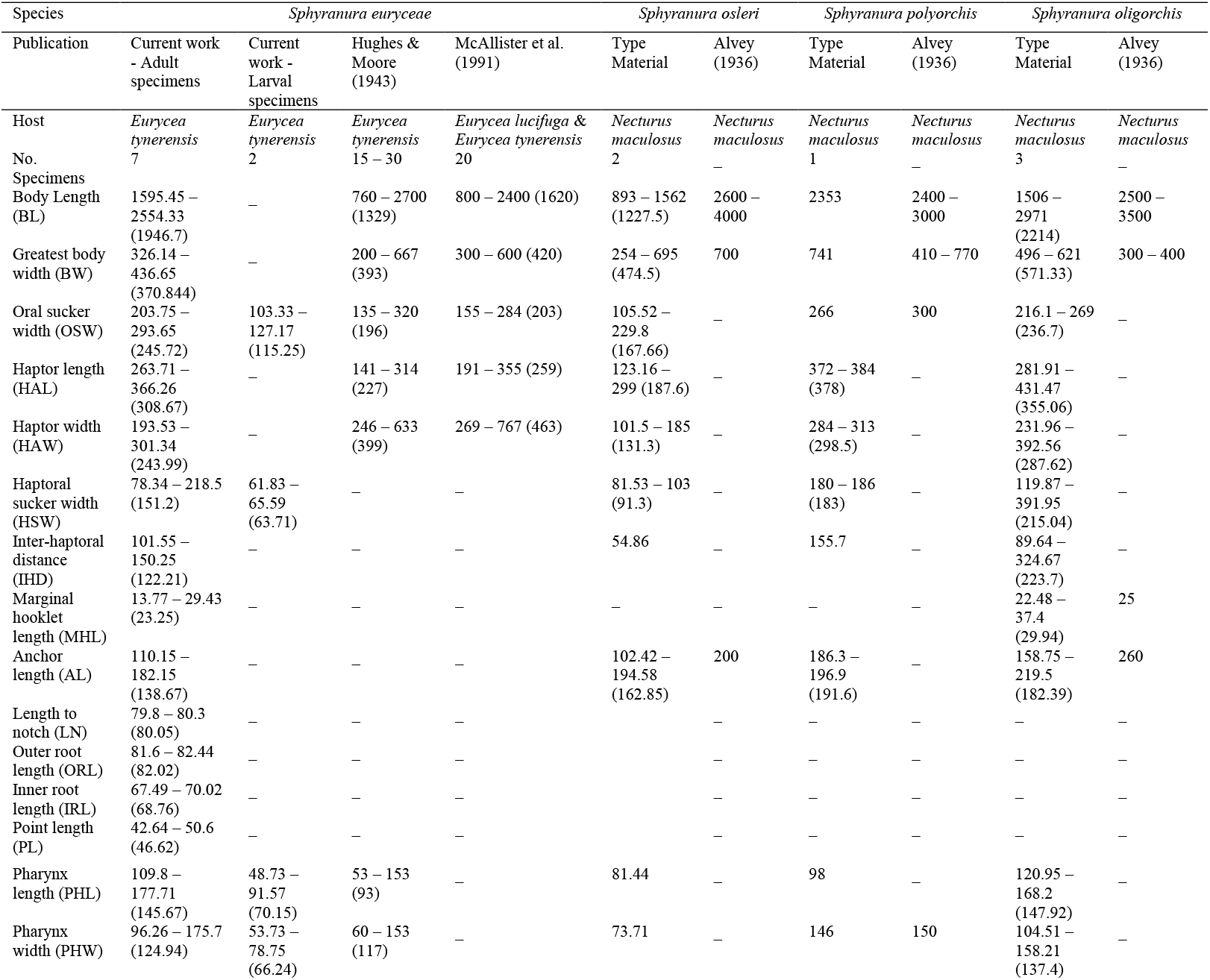
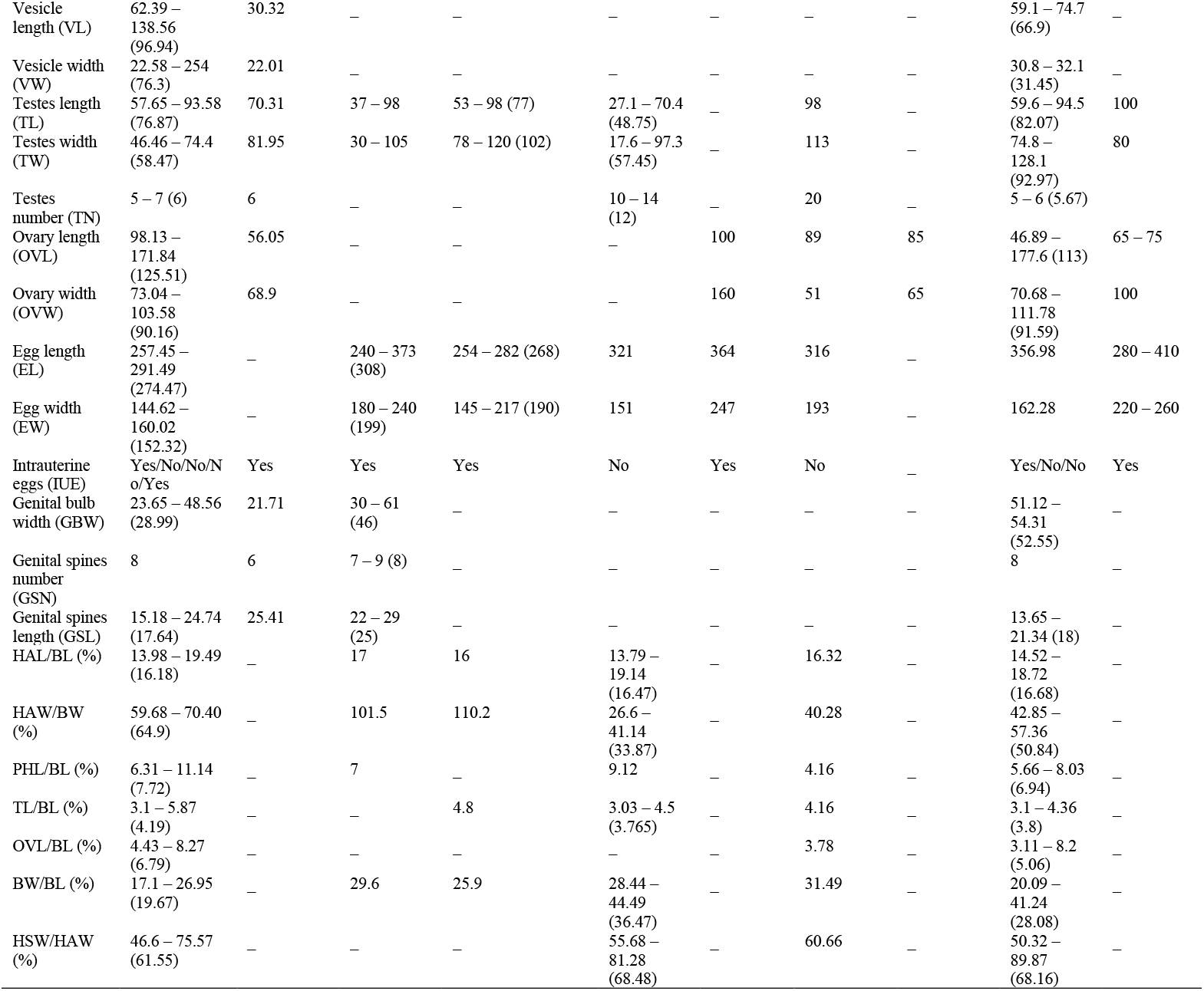
Morphological measurements in micrometres [µm] of new and previously published specimens of *S. euryceae* including re-measurement of type material of *S. osleri, S. oligorchis* and *S. polyorchis*. Range is followed by the mean in parentheses.

| Species | *Sphyrnura euryceae* |  |  |  | *Sphyrnura osleri* |  | *Sphyrnura polyorchis* |  | *Sphyrnura oligorchis* |  |
| --- | --- | --- | --- | --- | --- | --- | --- | --- | --- | --- |
| Publication | Current work - Adult specimens | Current work - Larval specimens | Hughes & Moore (1943) | McAllister et al. (1991) | Type Material | Alvey (1936) | Type Material | Alvey (1936) | Type Material | Alvey (1936) |
| Host | *Eurycea tynerensis* | *Eurycea tynerensis* | *Eurycea tynerensis* | *Eurycea lucifuga* & *Eurycea tynerensis* | *Necturus maculosus* | *Necturus maculosus* | *Necturus maculosus* | *Necturus maculosus* | *Necturus maculosus* | *Necturus maculosus* |
| No. Specimens | 7 | 2 | 15 – 30 | 20 | 2 | — | 1 | — | 3 | — |
| Body Length (BL) | 1595.45 – 2554.33 (1946.7) | — | 760 – 2700 (1329) | 800 – 2400 (1620) | 893 – 1562 (1227.5) | 2600 – 4000 | 2353 | 2400 – 3000 | 1506 – 2971 (2214) | 2500 – 3500 |
| Greatest body width (BW) | 326.14 – 436.65 (370.844) | — | 200 – 667 (393) | 300 – 600 (420) | 254 – 695 (474.5) | 700 | 741 | 410 – 770 | 496 – 621 (571.33) | 300 – 400 |
| Oral sucker width (OSW) | 203.75 – 293.65 (245.72) | 103.33 – 127.17 (115.25) | 135 – 320 (196) | 155 – 284 (203) | 105.52 – 229.8 (167.66) | — | 266 | 300 | 216.1 – 269 (236.7) | — |
| Haptor length (HAL) | 263.71 – 366.26 (308.67) | — | 141 – 314 (227) | 191 – 355 (259) | 123.16 – 299 (187.6) | — | 372 – 384 (378) | — | 281.91 – 431.47 (355.06) | — |
| Haptor width (HAW) | 193.53 – 301.34 (243.99) | — | 246 – 633 (399) | 269 – 767 (463) | 101.5 – 185 (131.3) | — | 284 – 313 (298.5) | — | 231.96 – 392.56 (287.62) | — |
| Haptoral sucker width (HSW) | 78.34 – 218.5 (151.2) | 61.83 – 65.59 (63.71) | — | — | 81.53 – 103 (91.3) | — | 180 – 186 (183) | — | 119.87 – 391.95 (215.04) | — |
| Inter-haptoral distance (IHD) | 101.55 – 150.25 (122.21) | — | — | — | 54.86 | — | 155.7 | — | 89.64 – 324.67 (223.7) | — |
| Marginal hooklet length (MHL) | 13.77 – 29.43 (23.25) | — | — | — | — | — | — | — | 22.48 – 37.4 (29.94) | 25 |
| Anchor length (AL) | 110.15 – 182.15 (138.67) | — | — | — | 102.42 – 194.58 (162.85) | 200 | 186.3 – 196.9 (191.6) | — | 158.75 – 219.5 (182.39) | 260 |
| Length to notch (LN) | 79.8 – 80.3 (80.05) | — | — | — | — | — | — | — | — | — |
| Outer root length (ORL) | 81.6 – 82.44 (82.02) | — | — | — | — | — | — | — | — | — |
| Inner root length (IRL) | 67.49 – 70.02 (68.76) | — | — | — | — | — | — | — | — | — |
| Point length (PL) | 42.64 – 50.6 (46.62) | — | — | — | — | — | — | — | — | — |
| Pharynx length (PHL) | 109.8 – 177.71 (145.67) | 48.73 – 91.57 (70.15) | 53 – 153 (93) | — | 81.44 | — | 98 | — | 120.95 – 168.2 (147.92) | — |
| Pharynx width (PHW) | 96.26 – 175.7 (124.94) | 53.73 – 78.75 (66.24) | 60 – 153 (117) | — | 73.71 | — | 146 | 150 | 104.51 – 158.21 (137.4) | — |
| Vesicle length (VL) | 62.39 – 138.56 (96.94) | 30.32 | – | – | – | – | – | – | 59.1 – 74.7 (66.9) | – |
| Vesicle width (VW) | 22.58 – 254 (76.3) | 22.01 | – | – | – | – | – | – | 30.8 – 32.1 (31.45) | – |
| Testes length (TL) | 57.65 – 93.58 (76.87) | 70.31 | 37 – 98 | 53 – 98 (77) | 27.1 – 70.4 (48.75) | – | 98 | – | 59.6 – 94.5 (82.07) | 100 |
| Testes width (TW) | 46.46 – 74.4 (58.47) | 81.95 | 30 – 105 | 78 – 120 (102) | 17.6 – 97.3 (57.45) | – | 113 | – | 74.8 – 128.1 (92.97) | 80 |
| Testes number (TN) | 5 – 7 (6) | 6 | – | – | 10 – 14 (12) | – | 20 | – | 5 – 6 (5.67) | – |
| Ovary length (OVL) | 98.13 – 171.84 (125.51) | 56.05 | – | – | – | 100 | 89 | 85 | 46.89 – 177.6 (113) | 65 – 75 |
| Ovary width (OVW) | 73.04 – 103.58 (90.16) | 68.9 | – | – | – | 160 | 51 | 65 | 70.68 – 111.78 (91.59) | 100 |
| Egg length (EL) | 257.45 – 291.49 (274.47) | – | 240 – 373 (308) | 254 – 282 (268) | 321 | 364 | 316 | – | 356.98 | 280 – 410 |
| Egg width (EW) | 144.62 – 160.02 (152.32) | – | 180 – 240 (199) | 145 – 217 (190) | 151 | 247 | 193 | – | 162.28 | 220 – 260 |
| Intrauterine eggs (IUE) | Yes/No/No/No/Yes | Yes | Yes | Yes | No | Yes | No | – | Yes/No/No | Yes |
| Genital bulb width (GBW) | 23.65 – 48.56 (28.99) | 21.71 | 30 – 61 (46) | – | – | – | – | – | 51.12 – 54.31 (52.55) | – |
| Genital spines number (GSN) | 8 | 6 | 7 – 9 (8) | – | – | – | – | – | 8 | – |
| Genital spines length (GSL) | 15.18 – 24.74 (17.64) | 25.41 | 22 – 29 (25) | – | – | – | – | – | 13.65 – 21.34 (18) | – |
| HAL/BL (%) | 13.98 – 19.49 (16.18) | – | 17 | 16 | 13.79 – 19.14 (16.47) | – | 16.32 | – | 14.52 – 18.72 (16.68) | – |
| HAW/BW (%) | 59.68 – 70.40 (64.9) | – | 101.5 | 110.2 | 26.6 – 41.14 (33.87) | – | 40.28 | – | 42.85 – 57.36 (50.84) | – |
| PHL/BL (%) | 6.31 – 11.14 (7.72) | – | 7 | – | 9.12 | – | 4.16 | – | 5.66 – 8.03 (6.94) | – |
| TL/BL (%) | 3.1 – 5.87 (4.19) | – | – | 4.8 | 3.03 – 4.5 (3.765) | – | 4.16 | – | 3.1 – 4.36 (3.8) | – |
| OVL/BL (%) | 4.43 – 8.27 (6.79) | – | – | – | – | – | 3.78 | – | 3.11 – 8.2 (5.06) | – |
| BW/BL (%) | 17.1 – 26.95 (19.67) | – | 29.6 | 25.9 | 28.44 – 44.49 (36.47) | – | 31.49 | – | 20.09 – 41.24 (28.08) | – |
| HSW/HAW (%) | 46.6 – 75.57 (61.55) | – | – | – | 55.68 – 81.28 (68.48) | – | 60.66 | – | 50.32 – 89.87 (68.16) | – |

***Sphyranura euryceae* Hughes & Moore, 1943**

**Type-host**: *Eurycea tynerensis* Moore & Hughes, 1939

**Other hosts**: *Eurycea lucifuga* Rafinesque, 1822 & *Eurycea spelaea* Stejneger, 1892

**Type-locality**: Pea Vine Creek, Cherokee County, Oklahoma, USA

**Other localities**: Greathouse Spring in Tontitown, Benton County, Arkansas, USA

**Type-specimens: Holotype:** US National Parasite Collection no. 36873 Hughes & Moore [18]. **Syntype:** USNM 1337573 Hughes & Moore [18]. **Vouchers:** USNM 1376383, McAllister [6], USNM 1398045 and 1398048 Bursey, USNM xx-xx present study, UH xx-xx present study

**Infection site:** Skin mainly at the base of legs, and external gills

**Infection parameters:** Current study - in 2019, 12 specimens of *Eurycea tynerensis* out of 27 infected (prevalence = 44,4%) with one or two individuals per host; in 2020, two out of six specimens of *E. tynerensis* infected (prevalence = 33,3%) with one individual. McAllister [6] reported infection in ten out of ten specimens of *E. lucifuga*, and ten out of ten specimens of *E. tynerensis* (prevalence = 100%). McAllister [17] reported infection in thirty-seven of seventy-four specimens of *E. tynerensis* and one of two specimens of *E. spelaea* (prevalence = 100%).

**Representative DNA sequences:** GenBank accession number xx-xx (*18S* rDNA), xx-xx (*28S* rDNA), xx-xx (*12S* rDNA), xx-xx (*cox1* mtDNA), xx-xx (mt genome)

#### Morphological Measurements

All measurements obtained in the course of the current study, both on new specimens and type material, as well as previous data on *Sphyranura* spp. is summarised in Table 2. The following measurements are reported as range followed by the mean in parentheses with all values reported in micrometres [µm]. Adult specimens of *S. euryceae* collected at Greathouse Spring (USA) for the current study are between 1595.45 – 2554.33 (1946.7) in length and 326.14 – 436.65 (370.844) at their greatest width. Oral sucker is subterminal and measured 203.75 – 293.65 (245.72) followed by the pharynx which is 109.8 – 177.71 (145.67) in length and 96.26 – 175.7 (124.94) in width. The ovary was observed in all adult specimens and measures 98.13 – 171.84 (125.51) in length and 73.04 – 103.58 (90.16) in width. Testes were observed in four of the seven adult specimens, numbering between 5 – 7 (6) per individual and measuring 57.65 – 93.58 (76.87) in length and 46.46 – 74.4 (58.47) in width. Haptors measured 263.71 – 366.26 (308.67) in length and 193.53 – 301.34 (243.99) in width with the haptoral sucker measuring 78.34 – 218.5 (151.2) in diameter. The distance between haptoral suckers is 101.55 – 150.25 (122.21). Haptors represent 13.98 – 19.49 (16.18) % of the body length and 59.68 – 70.40 (64.9) % of body width. The anchor exhibits an accessory sclerite at the base of the main hook and a deep, triangular cut between the inner 67.49 – 70.02 (68.76) and outer 81.6 – 82.44 (82.02) roots and a recurved hook with a point length of 42.64 – 50.6 (46.62). In addition to the seven adult specimens, morphological characteristics of two larvae were taken (Table 2). Micrographs showing morphological features of *S. euryceae* are presented in Figure 2.

**Figure 1.**
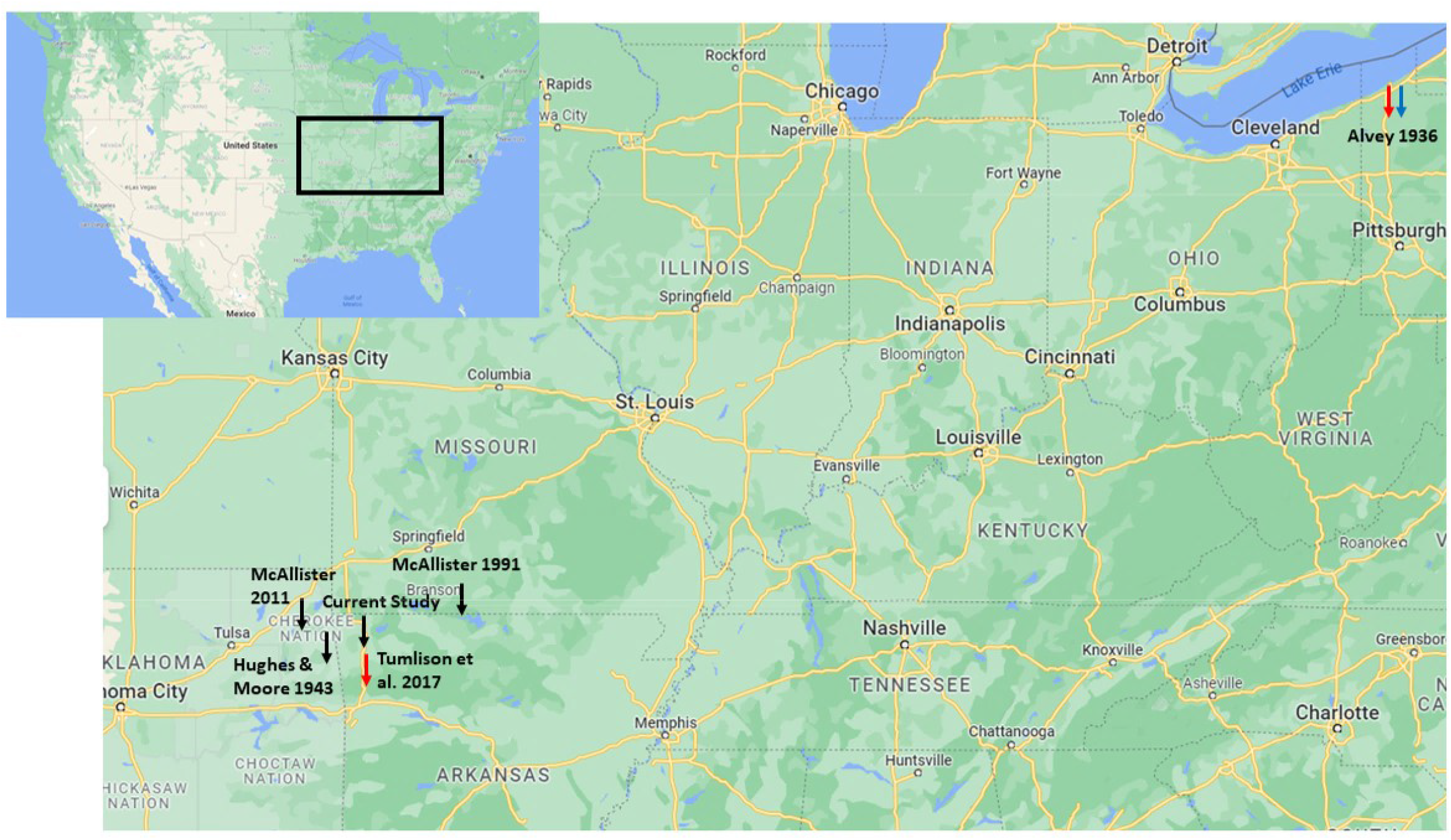
Geographic distribution of published records of *Sphyranura* where sampling location is available. Records of *S. euryceae, S. oligorchis* and *S. polyorchis* are marked in black, red, and blue, respectively.

**Figure 2.**
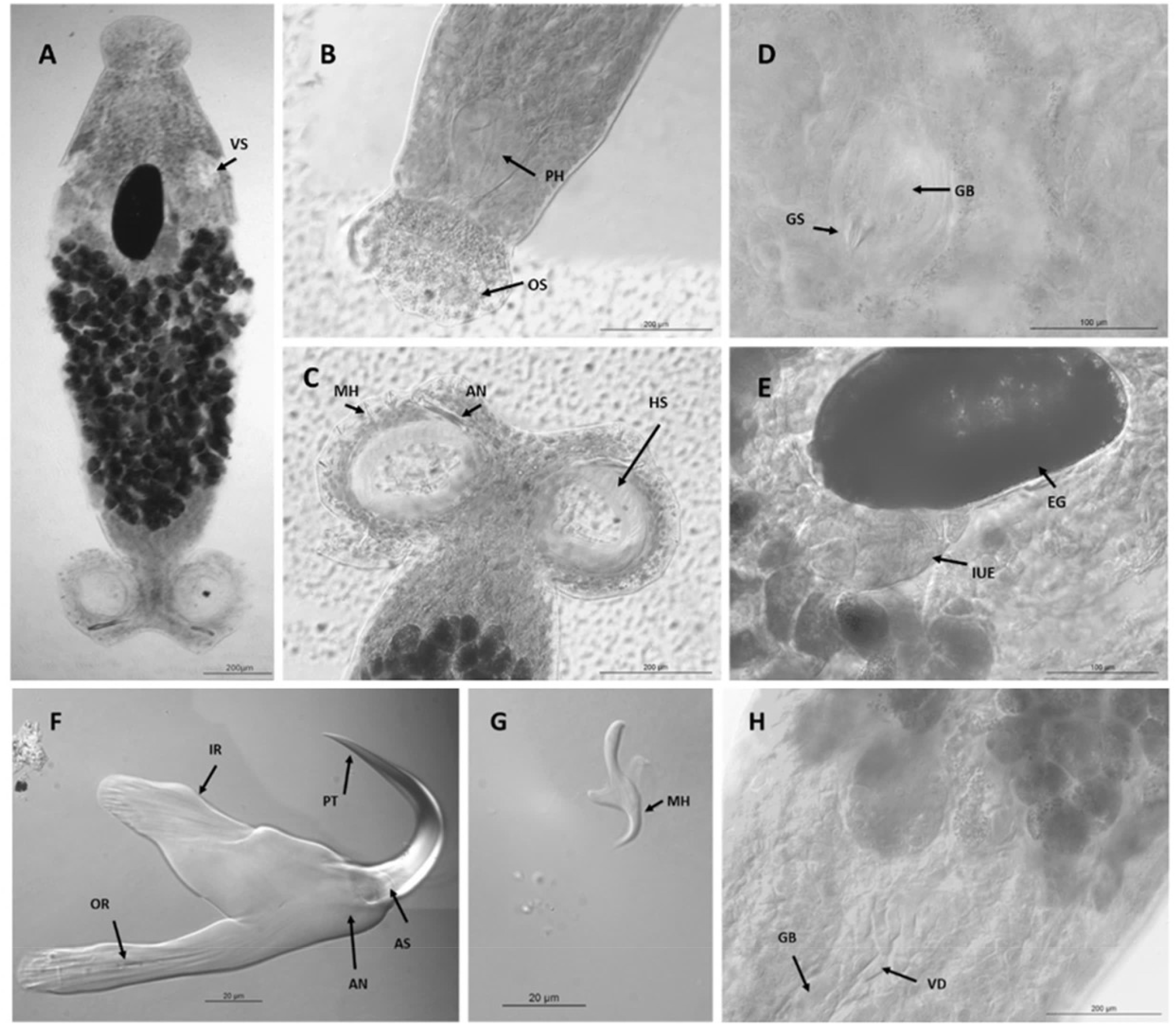
Micrographs of *Sphyranura euryceae*. A. Full body view, scale bar 200µm. B. Oral sucker and pharynx, scale bar 200µm. C. Haptor, scale bar 200µm. D. Genital bulb and spines, scale bar 100µm. E. Egg, scale bar 100µm. F. Anchor, scale bar 20µm. G. Marginal hooklet, scale bar 20µm. H. Vas deferens, scale bar 200µm. Abbreviations: PT, point, AN, Anchor, AS, accessory sclerite, IR, inner root, OR, outer root, MH, marginal hooklet, VS, vesicle, PH, pharynx, OS, oral sucker, GB, genital bulb, GS, genital spines, HS, haptoral sucker, EG, egg, IUE, intrauterine eggs, VD, vas deferens.

#### Differential diagnosis

*S. euryceae* may be distinguished from congeners on a number of morphological features. First, the overall body shape is more elongated than that of congeners (body width as a proportion of body length = 20% vs *S. osleri* = 36%, *S. polyorchis* = 31% and *S. oligorchis* = 28%) although there is some degree of overlap with *S. oligorchis*, but not with *S. osleri* and *S. polyorchis*. Further, haptor width as a proportion of body width is much greater in *S. euryceae* compared to the others (*S. euryceae* = 65% vs *S. osleri* = 34%, *S. polyorchis* = 40% and *S. oligorchis* = 51%). The oral sucker of *S. euryceae* is sub-terminal rather than terminal as in the other members of the genus. The mean anchor length of *S. euryceae* is also less than that of congeners although there is overlap between all species in this trait.

#### Mitochondrial genome

Mitochondrial genomes were assembled for the samples SPY1 and SPY2, a representation of which is presented in Figure 3. The assembly of SPY2 was performed using GetOrganelle from a subsample of 10 million reads, 41,406 of which were used post-filtering to assemble the mitochondrial genome. The assembly had a total length of 13,728bp and an average coverage of 201. Annotation of this assembly reveals the presence of 12 protein coding genes (the absence of atp8 is a characteristic of Neodermata [43]). Three non-coding regions with elevated AT content were found between *cox1* and *rrnL* (469bp, 78% AT), *nad6* and *nad5* genes (738 bp, 79% AT) and *cox2* and *cox3* genes (439 bp, 74% AT). A comparison of this mitochondrial genome with that of *D. hangzhouensis* is provided in Table 3. Overall, the two tRNA-genes missing in the original annotation of *D. hangzhouensis, trnV* and *trnA*, were found (see Table 5). Gene order differences of adjacent features between the two polystomatid species include *trnL2*/*trnS2* and *trnY*/*trnK*/*nad6*. Denovo assembly of SPY1 was attempted but did not successfully produce a full-length mitochondrial genome.

**Table 3.** Positions of mismatches between the sequences of SPY1 and SPY2 and the gene in which these are found.

| Position(s) | 946 | 1021 | 4812-4814 | 7235 | 8115 | 12040 | 12513 | 12918 |
| --- | --- | --- | --- | --- | --- | --- | --- | --- |
| SPY1 | A | A | TAA | T | G | G | C | G |
| SPY2 | T | T | - | C | A | A | T | T |
| Gene | *cox1* | *cox1* | *cox3* | *nad5* | *cob* | *trnA* | *nad1* | *nad1* |

**Figure 3.**
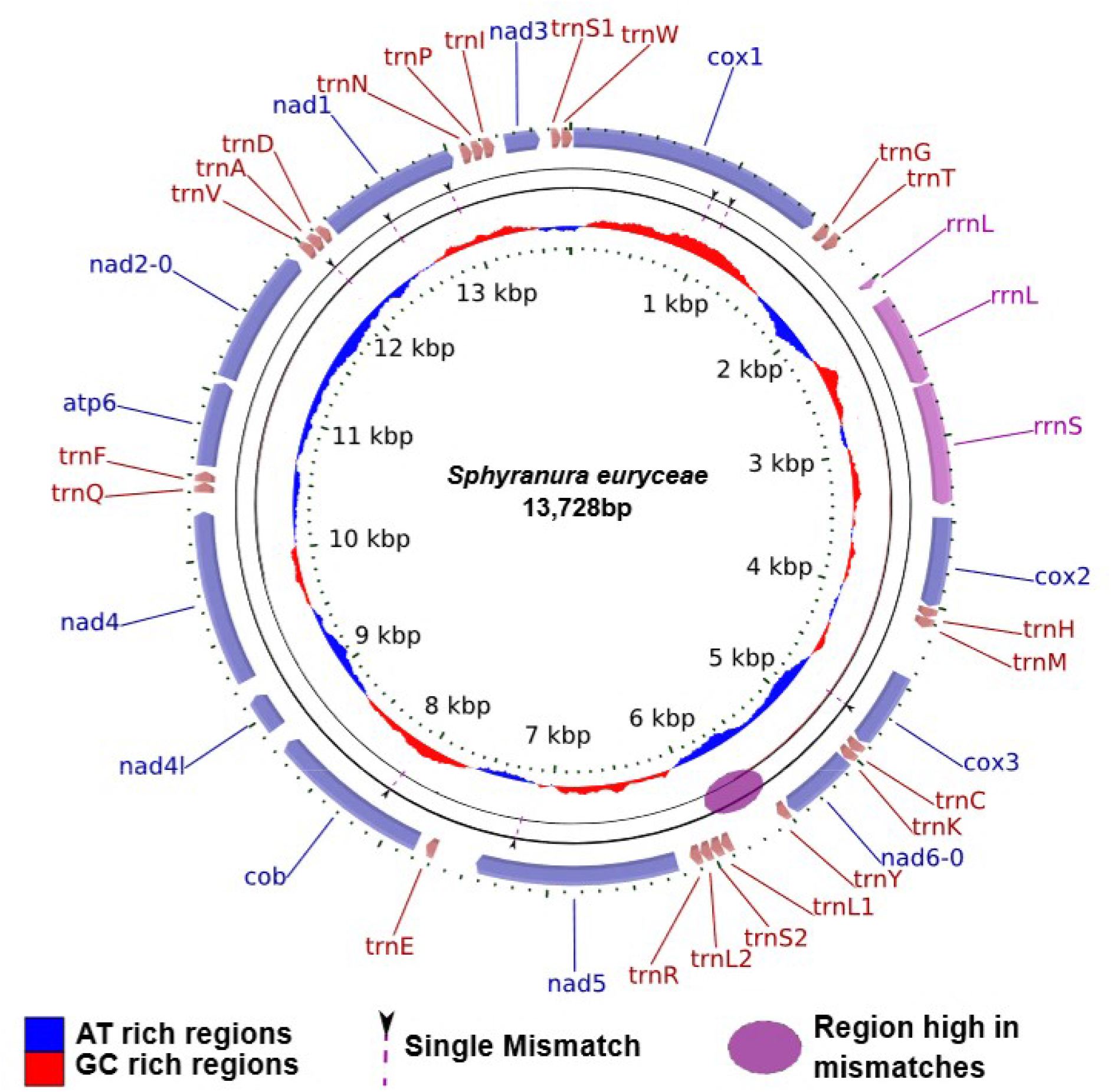
Visualisation of the annotated mitochondrial genome of *S. euryceae*. Mismatches between the samples SPY1 and SPY2 are indicated by dashed purple arrows and the region high in mismatches is indicated by the purple oval. AT rich regions are shown in blue in the inner circle whilst GC rich regions are shown in red.

However, when assembled using MITObim using the assembly of SPY2 as a reference, a full mitochondrial genome was recovered from a subsample of 10 million reads, 12,310 of which were mitochondrial. The two sequences were nearly identical with the following exceptions shown in Table 3. In addition to these differences there was a region of high dissimilarity between the positions 5545 and 5996. This dissimilarity was likely due to the presence of AT repeats which rendered this region difficult to assemble. Coverage differed between the two samples and is indicated in Table 4.

**Table 4.**
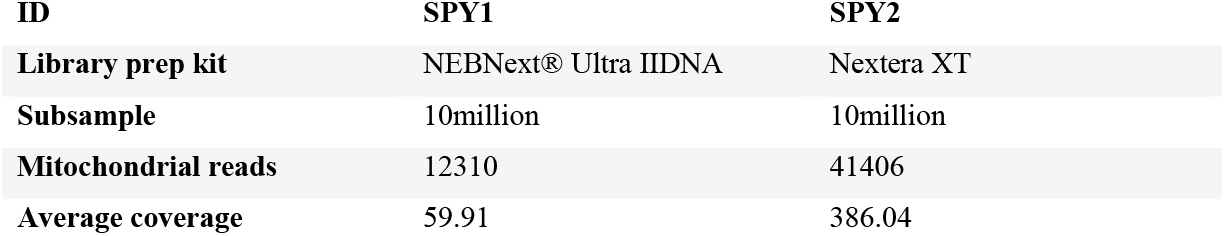
Library preparation kits and differing coverage of the sequences of SPY1 and SPY2.

**Table 5.**
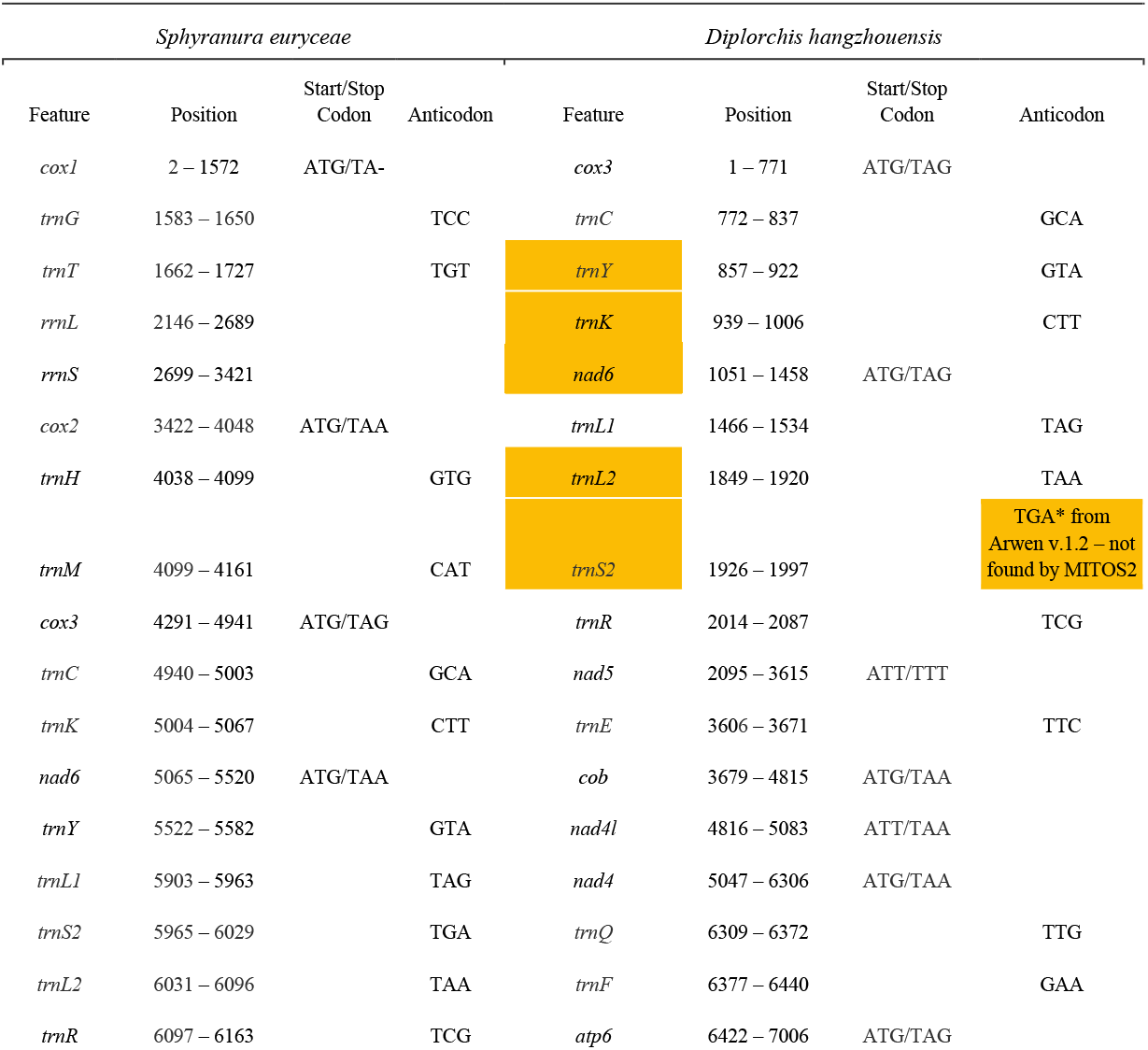
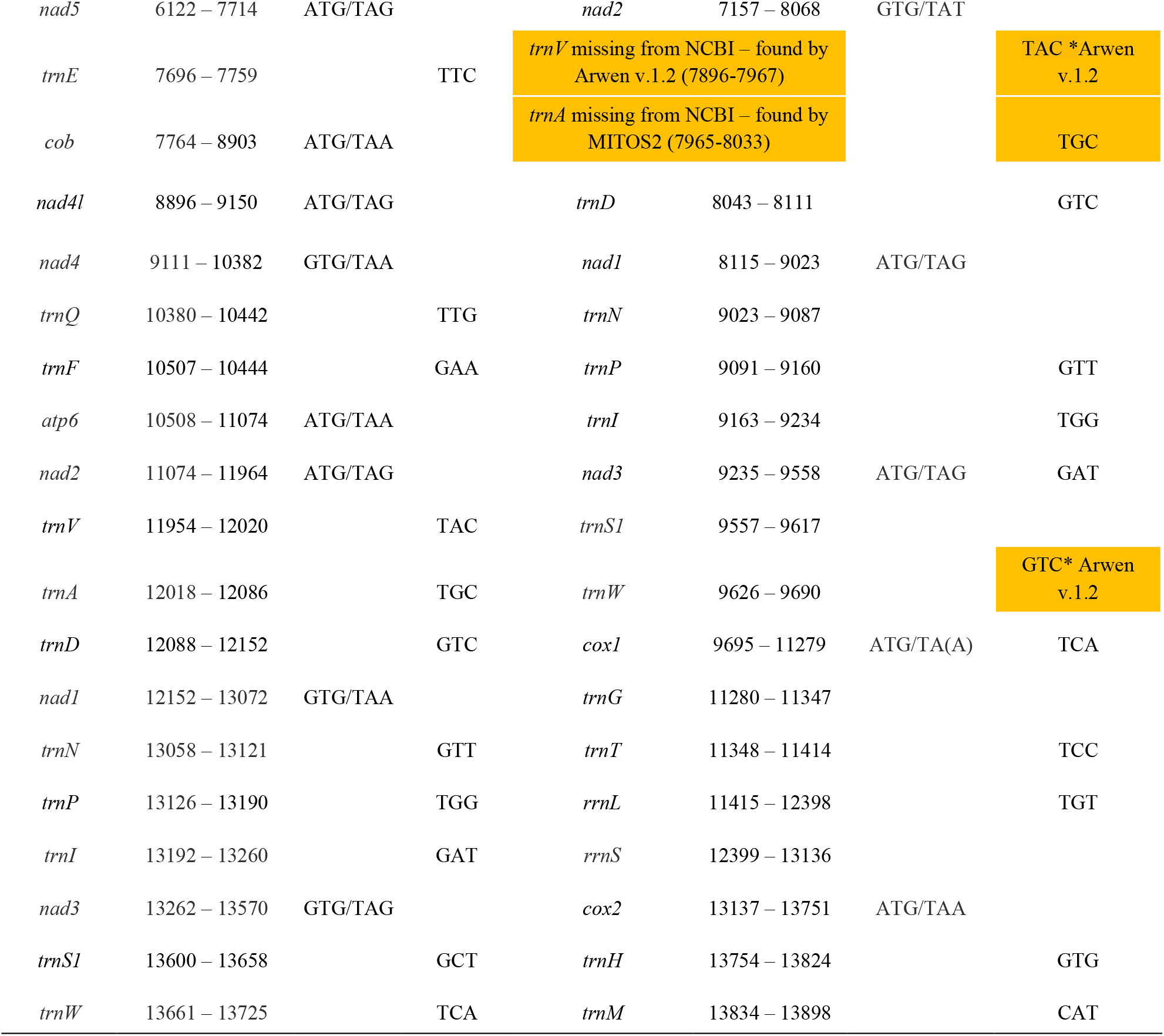
Comparison of mitochondrial genomes of *Sphyranura euryceae* and *Diplorchis hangzhouensis* (NCBI Accession JQ038227.1) including start and end positions of each feature, the start and stop codons of protein-coding genes and anticodons of tRNA genes. Instances in which gene order differs between the two species are highlighted in yellow. Positions given for *D. hangzhouensis* are as provided on NCBI. However, the *trnA* and *trnV* genes were not included on the NCBI annotation but were found in the present study, when reannotating the *D. hangzhouensis* with MITOS2.

#### Phylogenetic reconstruction

Sequences of *S. euryceae* were highly similar to those of *S. oligorchis* with percentage identities of 95.4% for *12S* (435 bp, including 15 indel positions), 99.1 – 99.2% for *18S* (2,009 bp), 100% for *28S* (1,411 bp) and 97.0 – 97.5% for *cox1* (395 bp). Maximum likelihood and Bayesian inference methods were employed on a total of 72 taxa (including 70 polystomatids and 2 non-polystomatid monogeneans) and resulted in phylogenies of Polystomatidae with broadly consistent topologies (Figure 4, Figure S2, Figure S3). These topologies were also largely in accordance with that inferred by Héritier et al. [21]. For instance, *C. australensis* was consistently resolved as a sister group of all other polystomatids. Furthermore, the two clades Héritier et al. [21] dubbed ‘Polbatrach’ and ‘Polchelon’ were also supported by the present study. In both Bayesian and maximum likelihood trees the new specimens of *Sphyranura euryceae* formed a monophyletic group that formed a sister-group relationship with *Sphyranura oligorchis*. Within the ‘Polbatrach’ clade *Sphyranura* emerges as the earliest branching lineage although this is weakly supported (0.75/43). Hence, the status of Sphyranuridae as a family separate to Polystomatidae is not supported by this topology. Inconsistencies were observed between the maximum likelihood and Bayesian trees at the next branch. In the maximum likelihood tree (Figure S2), we observe a split between the lineages leading on the one hand to the genera *Pseudodiplorchis, Neodiplorchis, Pseudopolystoma* and *Protopolystoma* and the genera *Diplorchis, Parapolystoma, Sundapolystoma, Polystoma, Madapolystoma, Kakana, Eupolystoma, Wetapolystoma* and *Metapolystoma* on the other. However, this node is not well supported [66]. The Bayesian tree (Figure 4, Figure S3) points to three lineages, containing *Protopolystoma, Pseudodiplorchis, Neodiplorchis*, and *Pseudopolystoma* branching independently, although also not well supported (0.71), prior to the well supported clade containing the genera *Diplorchis, Parapolystoma, Sundapolystoma, Polystoma, Madapolystoma, Kakana, Eupolystoma, Wetapolystoma* and *Metapolystoma*. In both ‘Polbatrach’ and ‘Polchelon’ clades many genera, including *Diplorchis, Polystoma, Neopolystoma* and *Polystomoides* appear non-monophyletic. There is further a general pattern of codivergence between host and parasite in the ‘Polbatrach’ clade, albeit with the exceptions of *P. dendriticum* and *Polystoma pelobatis* Euzet & Combes, 1966. However, such a pattern of codivergence is not seen in the ‘Polchelon’ lineage.

**Figure 4.**
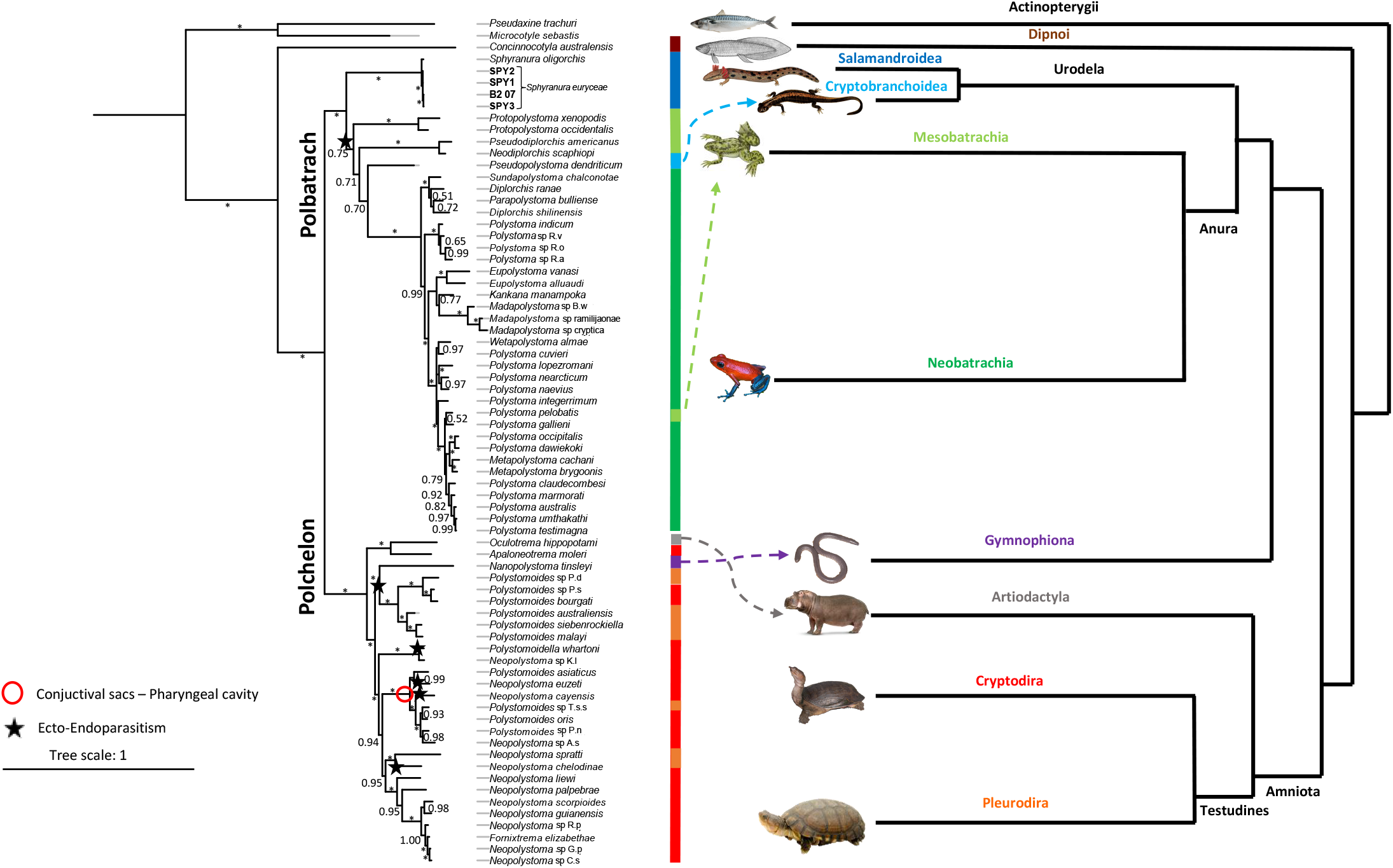
Bayesian Inference tree of Polystomatidae based on four concatenated nuclear (*18S* and *28S* rRNA) and mitochondrial (*12S* rRNA and *cox1*) gene portions. Schematic figure of hosts’ evolutionary history is based on [53]. Colours adjacent to taxon names indicate sub-order (in the case of Gymnophiona, order) of the host species. Stars indicate endoparasitic lineages. The red circle indicates a lineage parasitising the pharyngeal cavity. Tree scale bar represents 1 substitution per site.

## Discussion

*Sphyranura* was long thought to belong to Sphyranuridae. The first molecular phylogenies placed it at an early diverging, yet currently unresolved, position in the ‘Polbatrach’ clade of Polystomatidae. We provided an amended diagnosis of *Sphyranura* and obtained the first-ever molecular sequence data for *S. euryceae*. The inclusion of a second species of *Sphyranura* as well as 10 polystomatid taxa not included in previous phylogenies supports an early branching *Sphyranura* within the ‘Polbatrach’ clade and helped to elucidate the factors driving polystomatid phylogeny. Comparison at the mitochondrial genome level revealed instances of gene order differences in polystomatids.

### Morphological comparison of *Sphyranura* spp

Morphological analysis of the new specimens of *S. euryceae* and comparison of these with type material of *S. osleri, S. oligorchis* and *S. polyorchis* revealed high levels of both variability between conspecific individuals and overlap between each of the four species. It is important to note that individuals measured in this study as well as previous studies may well represent different life stages and may well have experienced different conditions prior to collection. Furthermore, the body tissues of monogeneans with the exception of the sclerotized attachment organs are soft and may not lie completely flat during the preparation of slides. For these reasons relative measurements should be used rather than absolute measurements for species differentiation. The most informative diagnostic features of *S. euryceae* however, included the following: an overall body shape which was elongated compared to congeners; greater haptoral sucker width in relation to body width; and a sub-terminal, rather than terminal oral sucker. Finally, anchor length of *S. euryceae* was also less than that of congeners. It should also be noted that type material measured in this study represented only a single individual of *S. polyorchis* of which many features were impossible to observe and measure. *S. osleri* was represented by two individuals, both deposited in 1879 and perhaps due to their age many features were again impossible to measure. Based on this, no definite conclusion should be drawn regarding the validity of *S. polyorchis* as questioned by Price [43].

### Mitochondrial Genome of *Sphyranura euryceae*

We provide the first available mitochondrial genome for *Sphyranura* and the second only for Polystomatidae. As with the majority of flatworm mitochondrial genomes available so far, 12 protein coding genes were found, with *atp8* being absent [44]. A further 21 tRNA genes and the genes coding for both the large and small subunits of the mitochondrial rRNA were present. Comparison with the mitochondrial genome of *D. hangzhouensis* reveals similar gene order, with two instances of rearrangement in the order of adjacent tRNA genes between the two species. However, the order of protein coding genes was conserved between the two species as has been observed in other monogenean families such as Dactylogyridae [45,46]. However, we identify differences in start/stop codon usage in 8 of 12 protein coding genes between the two polystomatids. Furthermore, the abbreviated stop codon (TA-) was used in *cox1* of *S. euryceae* whereas this stop codon was TAA in *D. hangzhouensis*. The fact that the mitochondrial genome of SPY1 could not be assembled de novo indicates that when performing library preparation with low input data the NEBNext® Ultra IIDNA Library Prep Kit is preferable to Nextera XT.

### Phylogenetic position of *Sphyranura*

The earliest branching lineage of Polystomatidae is *C. australensis* which parasitises the Australian lungfish. Given lungfish are the sister group to modern tetrapods [47], this indicates the evolution of Polystomatidae from fish to tetrapod parasites most likely occurred during the colonisation of terrestrial habitats by early tetrapods. The subsequent split between *‘*Polbatrach’ and ‘Polchelon’ clades further mirrors the divergence of Amphibia and reptilians. As first suggested by Sinnappah et al. (2001) and supported by Héritier et al. [21], *Sphyranura* is nested within the ‘Polbatrach’ clade of Polystomatidae, rendering Sphyranuridae invalid. *Sphyranura* therefore seems to represent a transitional state after the colonisation of tetrapods but prior to the shift from ectoparasitism to endoparasitism in the ‘Polbatrach’ clade that otherwise infects the urinary bladder. This is accompanied by the loss of two pairs of haptors in *Sphyranura*, a trait which Williams (1995) and Sinnappah et al. [19] attribute to paedomorphosis. As observed by Combes [48], Gallien [49] and Paul [48] polystomatid larvae first develop a single pair of haptoral suckers and so pass through a *Sphyranura*-like stage before developing the second and third haptoral sucker pairs. Gallien [49,50], and Paul [48] further observed that such larval stages may infect the external gills of anuran hosts in the tadpole stage thus accelerating their development to adults. It is therefore suggested that the colonisation of neotenic salamanders which retain a permanently aquatic lifestyle along with retention of external gills as adults resulted in the neotenic retention of an ectoparasitic, two-haptoral sucker state in adult members of *Sphyranura* [19]. The position of *Sphyranura* in our phylogeny as an early branching lineage in the ‘Polbatrach’ clade lends support to ectoparasitism in *Sphyranura* being an ancestral state whereas the endoparasitic lifestyle seen in later branching members of ‘Polbatrach’ is a derived state. However, the reduction in the number of haptoral suckers in *Sphyranura* can be understood as a derived character resulting from neotenic evolution. It is also of note that while the shift from ecto- to endoparasitism occurs once in the ‘Polbatrach’ clade, this shift occurred on five separate occasions in the ‘Polchelon’ clade. The transition from ectoparasitism of the conjunctival sacs to the pharyngeal cavity is attributed to a single event in the common ancestor of *Polystomoides asiaticus* Rohde, 1965, *P. oris* Paul, 1938, other species of *Polystomoides* and *Neopolystoma euzeti* Combes, 1976, *N. cayensis* Du Preez, Badets, Héritier & Verneau, 2017 and *Neopolystoma* sp. A.s. Furthermore, this may represent a transitional state between true ectoparasitism (such as on the eyes, skin or gills) and endoparasitism of the urinary bladder.

*Sphyranura* appears to form a sister group with all other members of ‘Polbatrach’ which occupy the urinary bladder of batrachians and possess six haptoral suckers. Apart from members of *Sphyranura, P. dendriticum* also parasitises a urodelan host, *Onychodactylus japonicus* Houttuyn, 1782 endemic to the Japanese islands of Honshu and Shikoku. Were a hypothesis of strict host-parasite phylogenetic congruence to hold true we should expect *P. dendriticum* to form a sister group with *Sphyranura*. Although both are members of the ‘Polbatrach’ clade, this is not supported by the data currently available. Instead, our phylogeny supports two independent acquisitions of urodelan hosts: one leading to *Sphyranura* in North America and another leading to *P. dendriticum* in Japan. Unlike the hosts of *Sphyranura, O. japonicus* goes through a full metamorphosis during which larvae lose their external gills [53]. As a result, the acquisition of caudatan hosts by the ancestor of *P. dendriticum* was accompanied neither by a shift to ectoparasitism nor a retention of larval morphology as seen in *Sphyranura*. It should further be noted that while both *N. maculosus* (the host of *S. osleri, S. oligorchis* and *S. polyorchis*) and members of the genus *Eurycea* (the hosts of *S. euryceae*) have a similar geographic distribution as well as neotenic retention of external gills as adults, they are not closely related, with *Necturus* belonging to Proteidae Gray, 1825 and *Eurycea* to Plethodontidae Wake, 1966. The evolutionary history of the ‘Polbatrach’ clade seems to mirror that of the host organisms at least at higher taxonomic levels. As discussed, our phylogeny points to *Sphyranura* as being the earliest branching lineage in the ‘Polbatrach’ clade with its members parasitising two families belonging to the sub-order Salamandroidea of Urodela. The branching order of the next three lineages is also not well supported but indicates divergence of *Protopolystoma*, parasitising Pipidae (belonging to Mesobatrachia), followed by the sister species *Pseudodiplorchis americanus* and *Neodiplorchis scaphopi* parasitising members of Scaphiopodidae (also belonging to Mesobatrachia) and finally *P. dendriticum*, a parasite of representatives of the sub-order Cryptobranchoidea of Caudata being an exception to the of co-divergent pattern. Next, we see the divergence of all remaining members of the ‘Polbatrach’ clade which parasitise representatives of various neobratrachian families with the exception of *Polystoma pelobatis*, which has undergone a host switch to members of Pelobatidae (belonging to Mesobatrachia). Therefore, with the exception of *P. dendriticum* and *P. pelobatis* which underwent host switches, we see a mirroring of the ‘Polbatrach’ phylogeny with that of Amphibia [54]. Such a pattern is not seen in the ‘Polchelon’ clade. Aside from the obvious exceptions of *Oculotrema hippopotami* and *Nanopolystoma tinsleyi* which respectively parasitise members of the non-chelonian orders of Artiodactyla Owen, 1848 and Gymnophiona Müller, 1832, there are also multiple switches between members of the chelonian sub-orders Cryptodira Cope, 1868 and Pleurodira Cope, 1864.

Geography does not seem to be a key driver of diversification patterns in Polystomatidae and does not explain the position of *Sphyranura* which is no more closely related to other North American polystomatids than members of ‘Polbatrach’ found on any other continent. There are many individual lineages in both the ‘Polbatrach’ and ‘Polchelon’ clades which are restricted to a given continent. This may be seen among other examples in *Protopolystoma* which is restricted to Africa, the sister species *P. americanus* and *N. scaphiopi* found in North America, the clade containing *Eupolystoma, Kankana and Madapolystoma* which is found in Africa with the branch leading to *Kankana* and *Madapolystoma* restricted specifically to Madagascar and the ‘Polchelon’ clade containing *Neopolystoma scorpioides* Du Preez, Badets, Héritier & Verneau, 2017, *N. guianensis* Du Preez, Badets, Héritier & Verneau, 2017, *Neopolystoma* spp. *and Fornixtrema elizabethae* Platt, 2000 found in the Americas. However, there are several cases in this family where sister species do not share such close geographic proximity. One stark example that cannot easily be explained by codivergence or geography is that of *N. tinsleyi* that parasitises the caecilian *Typhlonectes compressicauda* Duméril & Bibron, 1841. Héritier et al. [21] reported this species as the earliest branching of the ‘Polchelon’ clade and suggested that the colonisation of chelonians and subsequent diversification occurred once after the branching of *N. tinsleyi* and *O. hippopotami*. This sequence of events does not fit with our phylogeny, however, where the closest relatives of *N. tinsleyi* belong to the paraphyletic genus *Polystomoides*. Furthermore, *N. tinsleyi* is native to the amazon basin whereas its closest relatives are found in Africa, Australia, and South-East Asia. This topology therefore points to a colonisation of caecilians from an ancestor with a chelonian host likely prior to the splitting of the South American and African plates around 130 million years ago [55] given the limited dispersal abilities of these hosts.

The presence of not fully resolved nodes, particularly those leading to *Sphyranura, Protopolystoma, Neodiplorchis* and *Pseudodiplorchis* and *Pseudopolystoma*, leave outstanding questions regarding the final topology of Polystomatidae. Future efforts with the inclusion of further samples and potentially as yet undescribed species may resolve this. However, whilst there exist species of *Protopolystoma* for which molecular data are not yet present [56], *Neodiplorchis, Pseudodiplorchis* and *Pseudopolystoma* are monotypic at least as far as is understood from current literature. Additional sequences from polystomatid parasites of caecilians may also further our understanding of the phylogeny of Polystomatidae. While no such species are known besides *N. tinsleyi*, this species was only described in 2014. This coupled with the fact that caecilians are an understudied group does indicate that such undescribed parasite species may exist. However, more important than taxon sampling in resolving this phylogeny is access to more data, preferably on the genomic scale which unfortunately is currently unavailable.

## Conclusions

Our results support the conclusion that *Sphyranura* is nested within Polystomatidae, and that Sphyranuridae should be considered invalid. Furthermore, the apparent early branching position of *Sphyranura* indicates that its ectoparasitic lifestyle is most likely an ancestral trait with endoparasitism having evolved later in the ‘Polbatrach’ clade. On the other hand, the reduced number of haptoral suckers in *Sphyranura* compared to other polystomatids should be considered a derived rather than ancestral characteristic. This lends support to the hypothesis of Sinnappah et al. [19] that the ectoparasitic lifestyle and single pair of haptoral suckers of adult *Sphyranura* represents a case of an evolutionary retention of juvenile characters, given that other members of the ‘Polbatrach’ clade exhibit a two-haptor, ectoparasitic larval stage on the external gills of anuran tadpoles.

## Acknowledgements

This research was funded by Austrian Science Fund (FWF) (project P 32691). The Special Research Fund of Hasselt University supports M.P.M.V. (BOF20TT06) and N.K. (BOF21PD01). Specimens were collected under Scientific Collecting Permit (number 021120207) from the Arkansas Game and Fish Commission. Type material representing *Sphyranura oligorchis, S. osleri* and *S. polyorchis* was provided by the American Museum of Natural History.

## Supplementary Materials

**Figure S1**. *Sphyranura oligorchis* (AMNH1432.1). A. Full body view, scale bar 1000µm. B. Haptor, scale bar 100µm. C. Uterus and intrauterine eggs, scale bar 20µm. D. Pharynx, scale bar 20µm. E. Genital bulb and spines, scale bar 20µm. Abbreviations: PT, point, AN, Anchor, MH, marginal hooklet, V, vesicle, PH, pharynx, GB, genital bulb, GS, genital spines, HS, haptoral sucker, EG, egg, IUE, intrauterine eggs.

**Figure S2**. Maximum likelihood tree of Polystomatidae based on four concatenated nuclear (18S and 28S rRNA) and mitochondrial (12S rRNA and cox1) gene portions. Bootstrap values are indicated at the nodes. Those with support values lower than 80 are marked with *.

**Figure S3**. Bayesian Inference tree of Polystomatidae. Posterior probabilities are indicated at the nodes. Those with support values lower than 0.80 are marked with *.

