## Supplementary figures and images for "Ectoparasitism in Polystomatidae (Neodermata, Monogenea): phylogenetic position and mitogenome of *Sphyranura euryceae*, a parasite of the Oklahoma salamander"

### Fig.S1

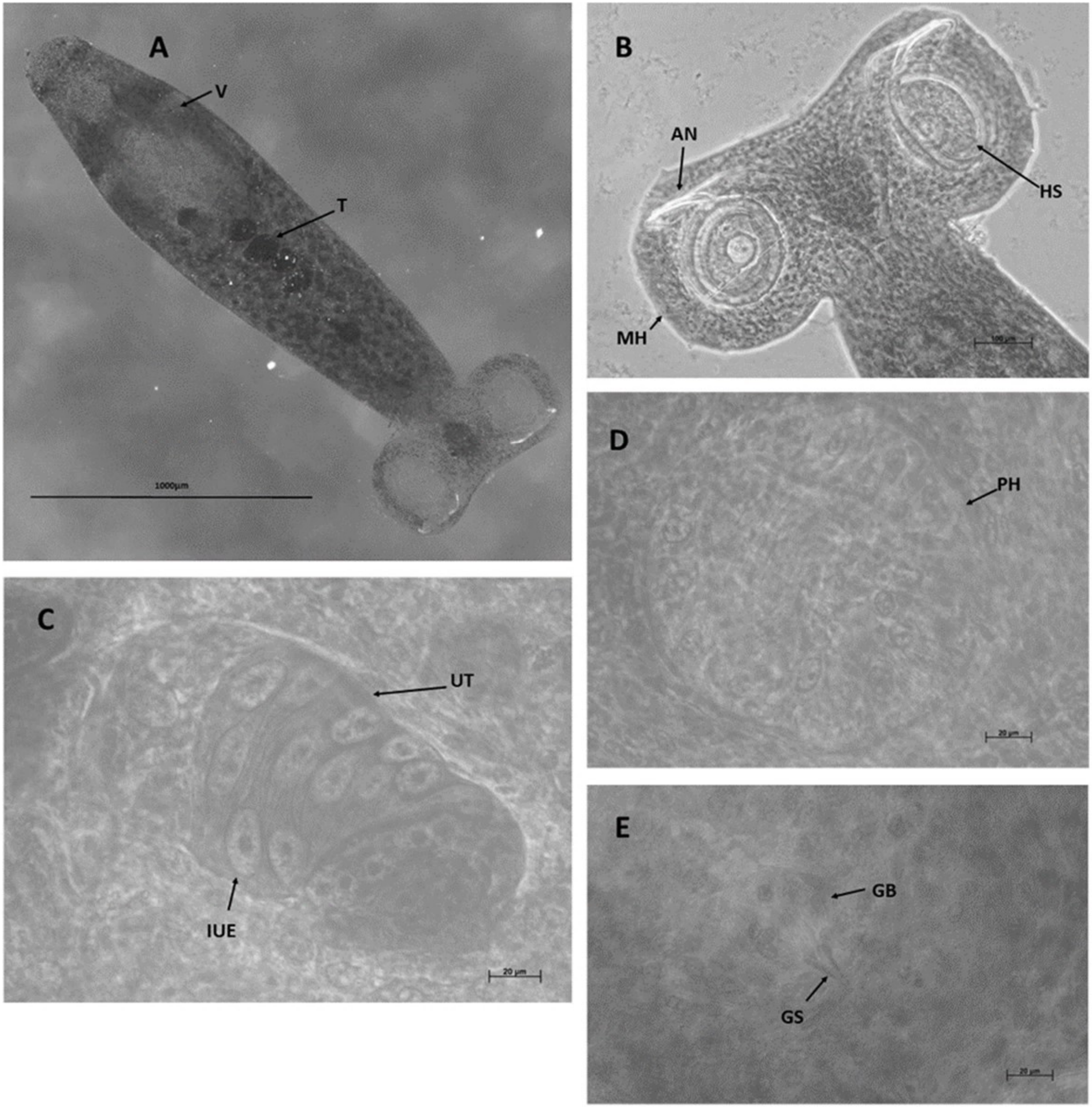

### Fig.S2

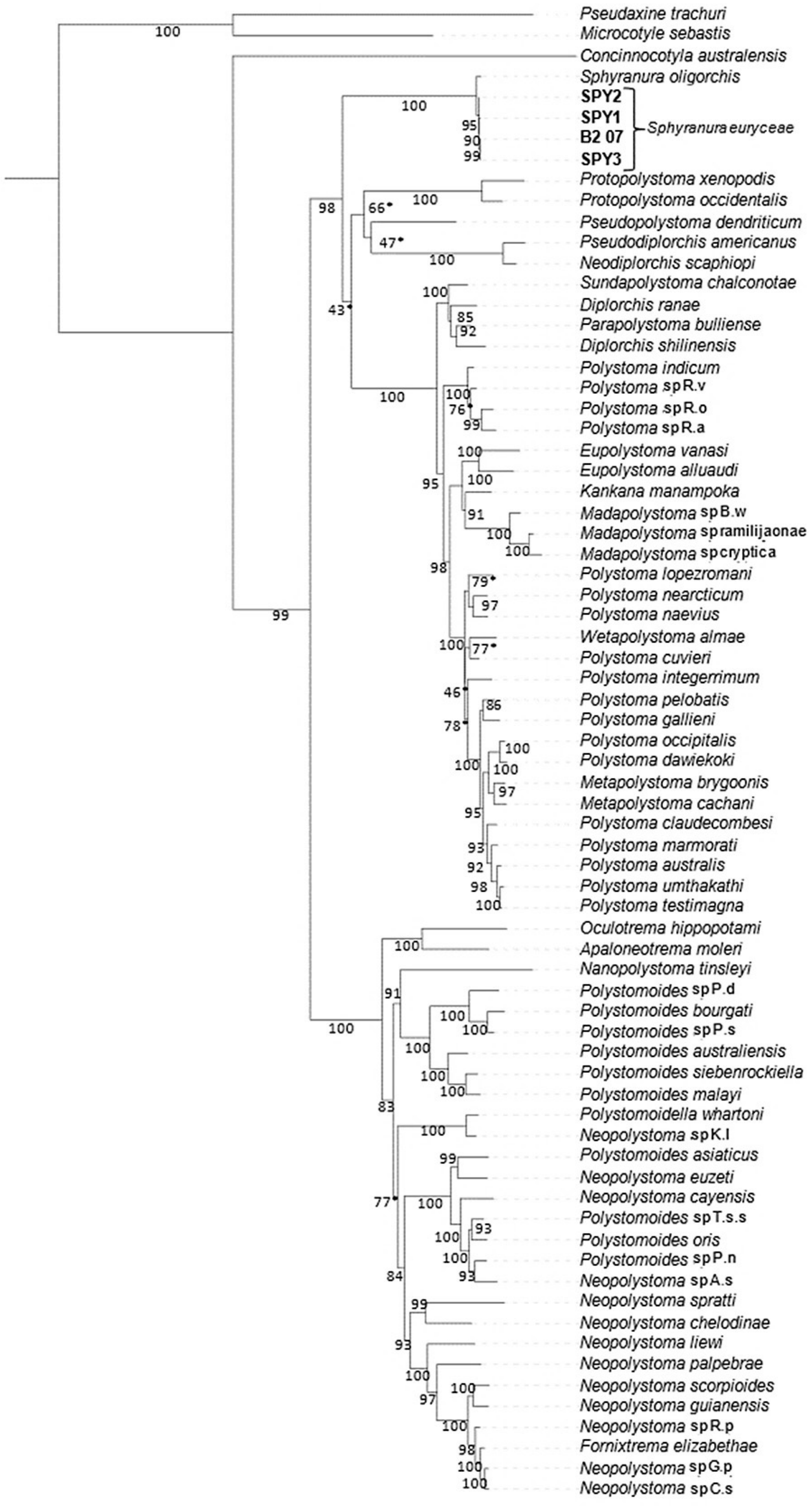

### Fig.S3

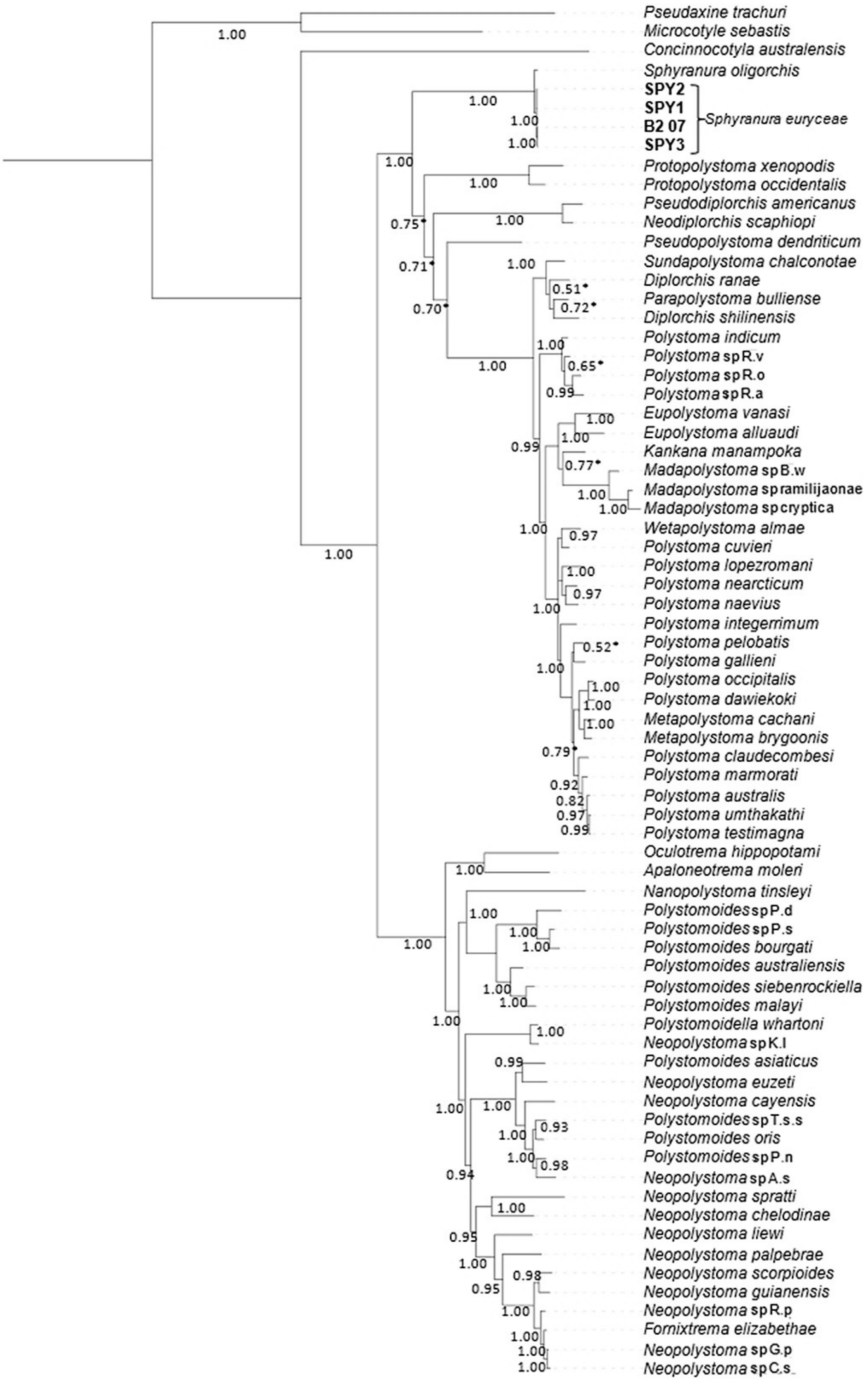
